# Ultra-sensitive water contaminant detection with transcription factor interfaced microcantilevers

**DOI:** 10.1101/2024.02.01.578376

**Authors:** Dilip K. Agarwal, Tyler J. Lucci, Jaeyoung K. Jung, Gajendra S. Shekhawat, Julius B. Lucks, Vinayak P. Dravid

## Abstract

Water contamination is a growing global concern, creating a need to develop technologies that can detect a range of target compounds at the required thresholds. Here, we address this need by merging biological allosteric transcription factors with DNA coated nanomechanical microcantilevers to detect chemicals in water with digital readout. After proof-of-concept demonstration and optimization to detect tetracycline with the TetR transcription factor, we use the CadC transcription factor to detect Pb^2+^ and Cd^2+^ in water at concentrations down to 2 ppb and 1 ppb, respectively, in less than fifteen minutes. A computational model suggests this improvement in sensitivity could be achieved by the DNA coated microcantilever surface changing transcription factor binding properties. Our findings demonstrate a promising new approach for water quality monitoring with fast, highly sensitive, digital readouts.

## INTRODUCTION

Reliable access to safe drinking water is a growing challenge, as studies estimate that 80% of the global population experiences high levels of threat to water security (*1*). One area of focus around water security is contamination of potable water supplies by harmful chemicals including ions, such as lead, cadmium, and fluoride, and small molecules, such as benzene, perfluoroalkyls, and glyphosate, which are toxic and can contribute to a range of negative health consequences (*2*). An ability to detect harmful chemicals in water quickly and accurately is thus critical to achieving water security for all.

Recent advances in synthetic biology have created cell-free biosensors that can detect harmful chemicals in water, offering an alternative to laboratory-based analytical chemistry approaches that are costly and slow (*3-6*). Cell-free biosensors work by repurposing allosteric transcription factors (aTFs) within cell-free gene expression reactions to sense chemicals in water (*3, 7-10*). For apo-repressor aTFs, reporter templates are configured to contain an aTF operator sequence in front of the reporter gene: in the absence of the target chemical, the aTF binds to the operator and blocks expression of the reporter; when the target chemical is present, it binds to the aTF, causing it to unbind from the operator and allow gene expression to continue (*3*). In this way, cell-free biosensors exploit the binding/unbinding of aTFs to DNA in the absence/presence of target chemicals to synthesize a reporter gene, which typically produces either a fluorescent (*3*) or colorimetric readout that can be detected by visual inspection.

Cell-free biosensing reactions can be assembled from biological components, lyophilized, and then used by simply adding a water sample and waiting to observe an output signal (*3*). By changing the aTF, they can be configured to detect a range of compounds including tetracycline (*3, 7*), Hg^2+^ (*7*), Cd^2+^ (*3*), Pb^2+^ (*3*), Azithromycin (*3*), and benzalkonium chloride (*3*), among others (*3*). Cell-free biosensors are also cost effective (<$1 USD/unit) (*3, 6*), can be manufactured and then distributed for use in the field at the point-of-need (*3, 6*), and can be operated by non-expert users (*11*), making them excellent candidates for increasing our collective capacity to scale water quality measurements to what is needed to address global challenges.

While promising, cell-free biosensors are still under development and currently have two key limitations. The first is with respect to limits of detection (LOD). Biosensor LODs are often determined by the biosensor system used. While for some contaminants such as fluoride, biosensors can achieve LODs that are relevant to EPA or WHO action limits (*6*), for others such as lead, biosensors have not been able to meet action limits (*3*). The second is readout quantitation. Most biosensors are configured to produce optical signals, making quantifying their signal difficult without sophisticated equipment not amenable to field deployment. There is thus great potential in interfacing cell-free biosensor reactions with digital readout platforms that can facilitate broad deployment, as recognized by recent work (*12*).

Here, we sought to address these limitations by creating an interface between cell-free biosensors and a highly sensitive nano-mechanical sensing platform that can be utilized to detect the fundamental binding/unbinding reactions that occur during aTF-mediated chemical detection. Specifically, we leverage microcantilevers – micrometer sized devices that physically bend when subjected to stresses induced by molecular binding events on their surfaces (*13-15*). When coated with binding interaction partners, they can be used to detect important molecules. For example, microcantilevers have been coated with antibodies, enabling them to detect a range of molecules through binding such as biotin (*15*), goat anti rabbit IgG (*15*), and SARS-CoV-2 spike protein (*16*), with high degrees of sensitivity (as low as hundreds of fM (*16*)). In addition to detection sensitivity, microcantilevers have a number of advantages including their small size (500 x 95 x 1 μm (*16*)), ability to parallelize in a detection array (*17, 18*), low sample volume requirements (as low as 4 μL (*14*)), and their ability to provide a continuous digital readout (*15*). When coupled with MOSFET technology, microcantilevers are even compatible with direct monolithic integration for integrated circuits, enabling their deployment in a wide range of electronic devices (*15*).

To address current limitations of cell-free biosensors and expand the capabilities of nano-mechanical sensors, we created a new approach to interface the RNA Output Sensors Activated by Ligand INDuction (ROSALIND) (*3*) aTF biosensing system with microcantilevers by patterning the microcantilever surface with DNA containing aTF operators (**Fig. 1**). In this way, binding of the aTF to the DNA causes microcantilever bending, while the presence of a specific chemical that releases the aTF from the DNA causes microcantilever de-bending. We first established the system using optical detection of microcantilever deflection and DNA encoding the operator sequence for the TetR aTF which senses tetracycline. After exploring the effects of incorporating multiple aTF operators within the bound DNA, we then configured the system to detect lead and cadmium with the CadC aTF. Using this approach, we show the ability to detect lead and cadmium in water at concentrations down to 10 nM each (2 ppb for lead and 1 ppb for cadmium). Notably, these levels are below the EPA practical quantitation level of 5 ppb for lead (*19*) and EPA maximum contaminant level goal (MCLG) of 5 ppb for cadmium (*2*), and improve upon those demonstrated by ROSALIND by approximately two orders of magnitude (*3*). In addition, we show that these results can be obtained within 15 minutes (sample to answer ≤ 15 min) with a digital readout. We anticipate these results will open the door to new approaches for rapid detection of water contaminants with platforms that can be embedded in portable digital devices.

**Fig. 1.**
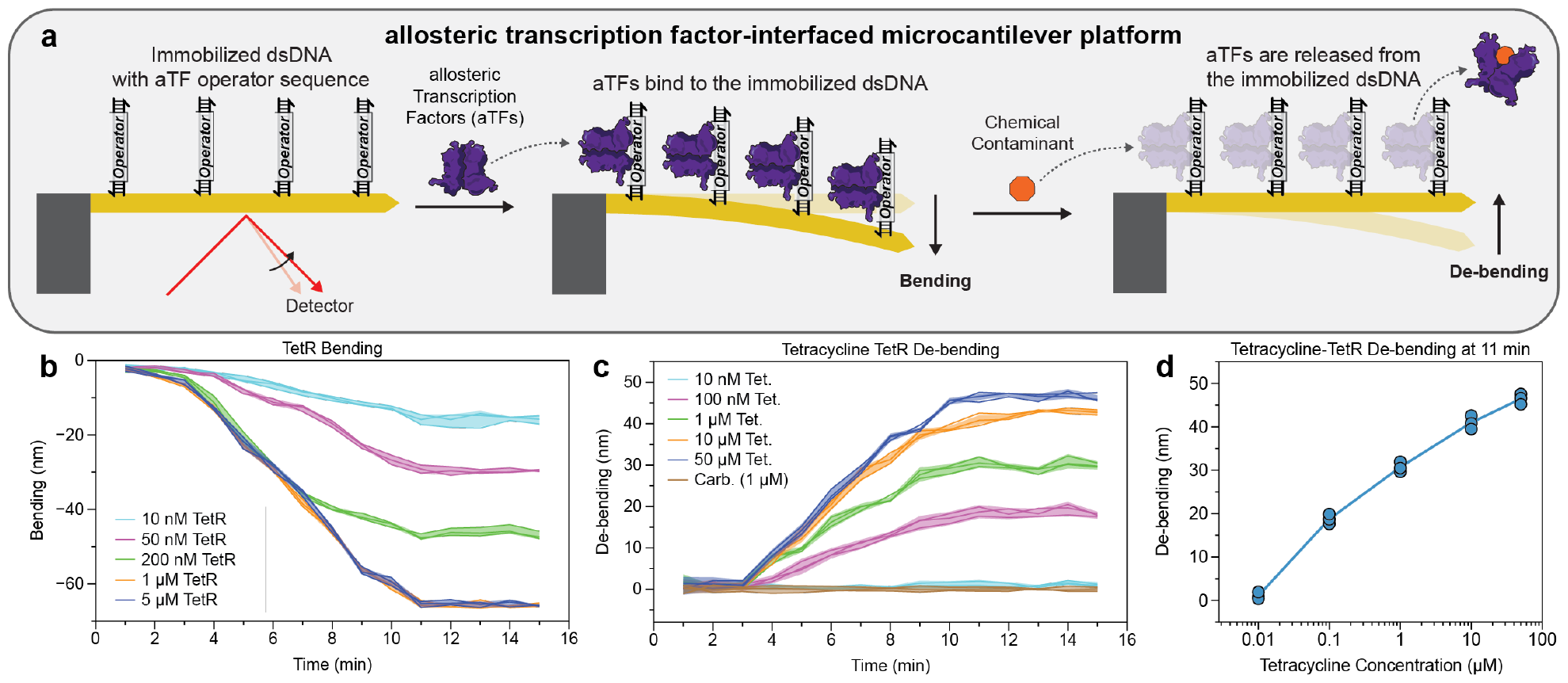
Interfacing microcantilevers with allosteric transcription factor biosensors. a) Microcantilevers are first coated with double stranded DNA (dsDNA) containing aTF operator sequences. aTF binding to the immobilized dsDNA causes cantilever bending. Binding of a specific chemical contaminant to the aTF causes an allosteric change, releasing aTFs from the dsDNA, resulting in cantilever de-bending which can be detected optically. b-d) Interfacing a microcantilever with the aTF TetR which binds to a DNA operator sequence, *tetO*, and responds to the antibiotic chemical tetracycline. b) Microcantilever bending in response to various concentrations of TetR. c) Microcantilever de-bending with 1 μM TetR and various concentrations of tetracycline. Carbenicillin, an antibiotic not recognized by TetR, was used as a control. Debending is computed as the difference in bending (deflection) between the sensing microcantilever, induced with tetracycline, and the average bending of the control microcantilevers, induced with 1 μM carbenicillin (**Fig. S2**). d) A dose response curve calculated from the kinetic traces in c) after 11 minutes. The solid line passes through the mean of the data points. Data in b-d are for n = 3 technical replicates, where each technical replicate is represented by a solid line (b, c) or point (d). Shaded regions in b, c represent plus/minus one standard deviation about the mean of all technical replicates at each condition. Full data plots can be found in **Fig. S3**, and full data can be found in the **Supplementary Data**.

## RESULTS

### DNA patterned microcantilevers can detect transcription factor binding and unbinding in response to external ligands

Our first goal was to test whether microcantilevers could detect transcription factor binding via patterning their surface with DNA containing aTF operator sequences (**Fig. 1a**). To assemble this system, we used a similar approach to others who have demonstrated protein detection with DNA-functionalized microcantilevers (*20*). Here, gold-coated microcantilevers were functionalized by immersing them in a solution of thiolated double stranded DNA (dsDNA) containing operator sites corresponding to a specific aTF. As a proof-of-concept, we began with the well-characterized tetracycline-TetR repressor system (*21*) and included a single *tetO* operator within the 19 bp dsDNA sequence (**Fig. 1a**).

DNA-coated microcantilevers were next placed inside a microfluidic chip (**Fig. S1**), and 10 μL of solution containing TetR at the specified concentration was added to the chip via pipette. Microcantilever bending was then detected optically using an AFM instrument as a proof-of-concept (see Methods). Exposing the microcantilever to a range of TetR concentrations (10 nM to 1 μM) increased bending magnitude over time, stabilizing within approximately 11 minutes of TetR addition for all TetR concentrations studied (**Fig. 1b**). In addition, the stabilized bending magnitude increased with increasing TetR, up to a concentration of 1 μM, after which it saturated (**Fig. 1b**). We therefore chose to proceed with the 1 μM TetR condition.

We next sought to investigate whether we could observe a change in microcantilever bending when the system is exposed to the TetR ligand, anhydrotetracycline (tet) (*21*). To test this, DNA-coated microcantilevers were exposed to 10 μL of 1 μM TetR for a period of approximately 15 min to allow complete bending, and then subjected to 10 μL of a solution containing either tetracycline or the ligand control carbenicillin, which is known to not interact with TetR (*22*). Notably, we did not observe a change in bending due to the addition of carbenicillin, indicating that TetR binding is not affected by the change in solution volume during this procedure. This allowed us to quantify the decrease in bending, denoted as ‘de-bending’, as the difference between the cantilever deflection in the presence of carbenicillin from that observed with a particular tetracycline condition at each point in time (see **Methods, Fig S2**). Using this approach, we observed an increase in de-bending over time, stabilizing within approximately 11 minutes of tetracycline addition (**Fig. 1c**). In addition, statistical analysis of de-bending trajectories showed that there are significant differences in de-bending magnitude above baseline at 100 nM anhydrotetracycline under 5 min. Plotting the de-bending magnitude at 11 minutes versus anhydrotetracycline concentration showed an increase over the concentration range of 10 nM to 100 μM (**Fig. 1d)**.

Overall, this data demonstrated the ability of the DNA-coated microcantilever to detect the binding of aTFs, and their release upon exposure to cognate ligands within several minutes, demonstrating the proof of concept of our approach and the speed of our system.

### Varying DNA operator number and spacing enhances detection sensitivity

Next, we sought to understand if modifying the DNA design by adding multiple operator sites per dsDNA and altering the spacing between the operator sites would influence sensitivity. For the tetracycline-TetR system, we investigated one operator site per dsDNA (“1X”), two operator sites per dsDNA with three base pair spacing between them (“2X3BP”), two operator sites per dsDNA with six base pair spacing between them (“2X6BP”), and three operator sites per dsDNA with six base pair spacing between them (“3X6BP”) (**Fig. 2a**).

**Fig. 2.**
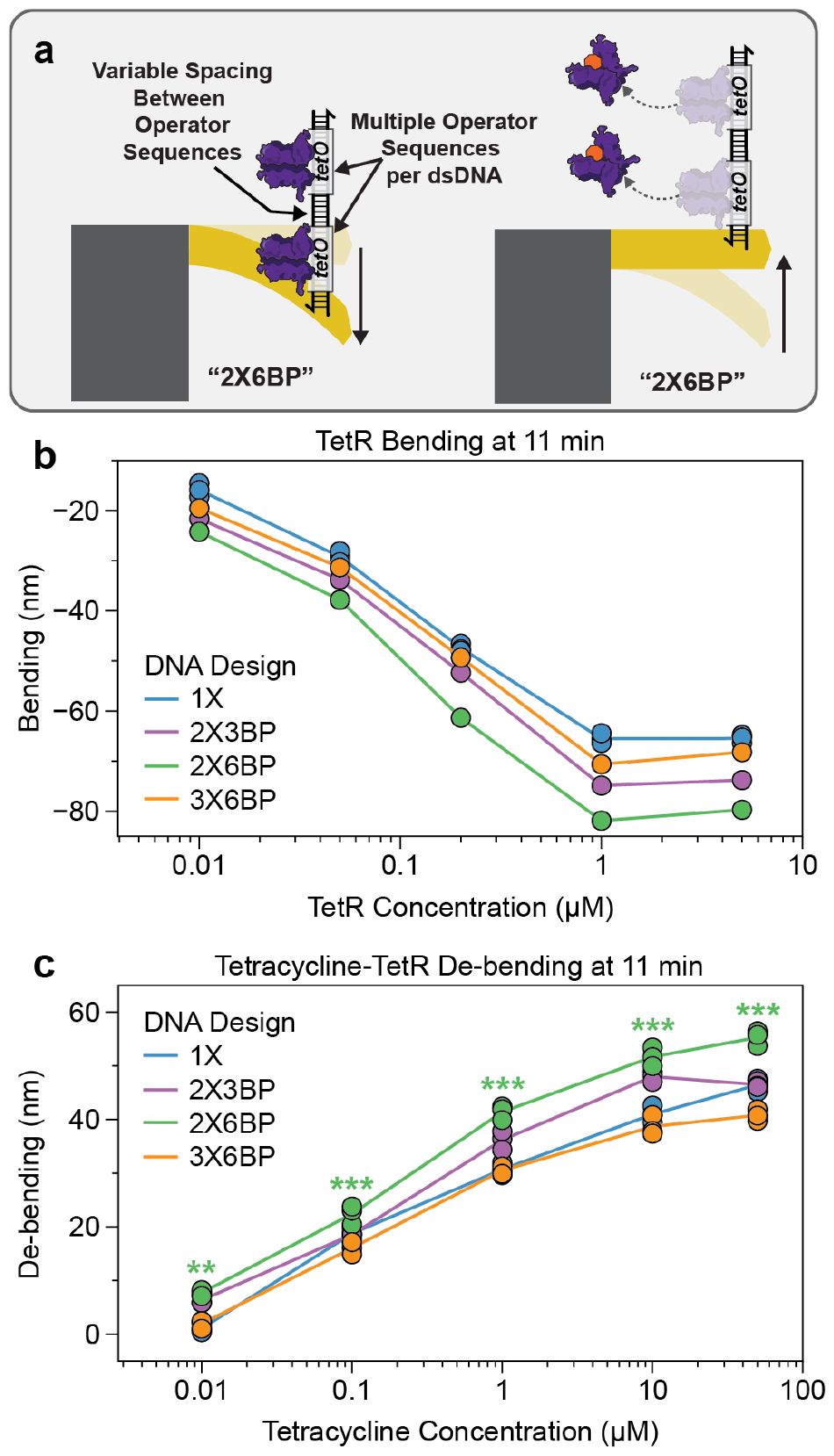
Varying the number and spacing of DNA operators can tune microcantilever detection performance. a) DNA sequences were modified to contain multiple TetR operator sites with different spacings between them. The notation NXMBP indicates N operator sites were included with a spacing of M bps. b) Bending data collected at 11 minutes with different DNA designs for the TetR system over a range of TetR concentrations. c) De-bending data collected at 11 minutes using 1 μM TetR and a range of tetracycline with different DNA designs. De-bending is calculated using a 1 μM carbenicillin control condition as in **Fig. 1**. Individual data points represent technical replicates, each corresponding to a single microcantilever. Solid lines in b) connect data points while in c) pass through the mean of the data points. For the 2X6BP case, the ligand concentration at which the bending magnitude was distinguishable from the carbenicillin induction control were determined using a two-tailed, homoscedastic Student’s t-test against the carbenicillin control condition, and their *P* value ranges are indicated with asterisks (\*\*\**P* < 0.001, \*\**P* = 0.001–0.01, \**P* = 0.01–0.05). Time course data plots can be found in **Fig. S3**, and full data and P values can be found in the **Supplementary Data**.

Microcantilevers with different DNA designs were prepared and used for both bending and de-bending reactions as before. Increasing the number of operator sites from one to two caused a greater magnitude of bending with a given TetR concentration, which was further increased by increasing the spacing of the two operator sequences from three to six bp (**Fig. 2b**). This translated into an increased sensitivity of tetracycline detection in de-bending experiments, with the 2X6BP design showing consistently larger levels of de-bending for tetracycline levels above 10 nM (**Fig. 2c**). Performing Student’s t-tests between the time course trajectories of the tetracycline and carbenicillin controls at 11 minutes revealed that the 2X6BP design showed statistically significant differences at 10 nM tetracycline. This allowed us to define the limit of detection (LOD) for the 2X6BP system at 10 nM tetracycline, which is notably lower than that observed for the previously characterized ROSALIND transcription system of ∼1 μM when using TetR to control the expression of a fluorescent RNA aptamer for detection, and ∼100 nM when using a negative feedback genetic circuit (*3*). Interestingly, further improvements in bending and de-bending sensitivity were not observed for the 3X6BP design, suggesting that there is a limit to this approach of tuning.

These results demonstrate that DNA design is an additional handle with which the aTF-microcantilever system can be tuned, and that the system has improved sensitivity over cell-free gene expression systems that utilize the same aTFs.

### Single digit parts per billion detection of lead and cadmium can be achieved with microcantilevers interfaced with the CadC transcription factor

We next sought to extend our sensor platform to the detection of heavy metals that can lead to serious health consequences if present in drinking water (*23, 24*). Lead is of particular concern, and lead concentration is tightly regulated by the US EPAs Lead and Copper Rule (LCR) (*19*). The LCR sets an action level of 15 parts per billion (ppb, 72 nM) for lead and stipulates that water systems must take action via public education or lead service line replacement if 10% of the samples collected from households using that water source exceed this level (*25*).

To configure the microcantilever system to detect lead, we utilized the well-characterized transcription factor CadC from *Staphylococcus aureus* (*26, 27*), which has been previously used in the ROSALIND system (*3*). CadC also has the advantage of being able to detect cadmium (*27, 28*) and does so through a similar mechanism as TetR, except it recognizes a different 22 bp DNA operator sequence *cadO* (*29*). We replaced *tetO* with the *cadO* operator sequence in the 2X6BP DNA template design (**Fig. 2**) and characterized bending with respect to the addition of CadC and either lead or cadmium (**Fig. S4, S5**). Similar to microcantilever bending with TetR (**Fig. 1a**), our results show that microcantilever bending magnitude increases with increasing CadC concentration from 10 nM to 5 μM after which it plateaus at higher concentrations (**Fig. S4, S5**). In addition, the bending magnitude stabilizes within approximately 11 minutes of CadC addition for all CadC concentrations studied. The optimized concentration of 5 μM CadC was therefore chosen to obtain maximum microcantilever bending prior to ligand (lead or cadmium) addition.

We next studied de-bending when subjecting CadC-bound microcantilevers to either lead or cadmium (**Fig. 3b-c**). In both cases, we were able to observe an increasing de-bending response when subjected to increasing concentrations of lead and cadmium. Using the same analysis to determine LOD, we found an LOD at 11 minutes of 10 nM (2 ppb) for lead and 10 nM (1 ppb) for cadmium, both lower than the EPA action limits of 15 ppb and 5 ppb for lead and cadmium (*2, 19*), respectively. Notably these values were two orders of magnitude lower than those demonstrated previously with ROSALIND, which were 1.2 μM (250 ppb) for lead and 1.25 μM (140 ppb) for cadmium (*3*). Interestingly, we did not observe a distinguishable de-bending response from exposure to 50 μM of Zn^2+^ compared to the carbenicillin control for the CadC system (**Fig. S4**).

**Fig. 3.**
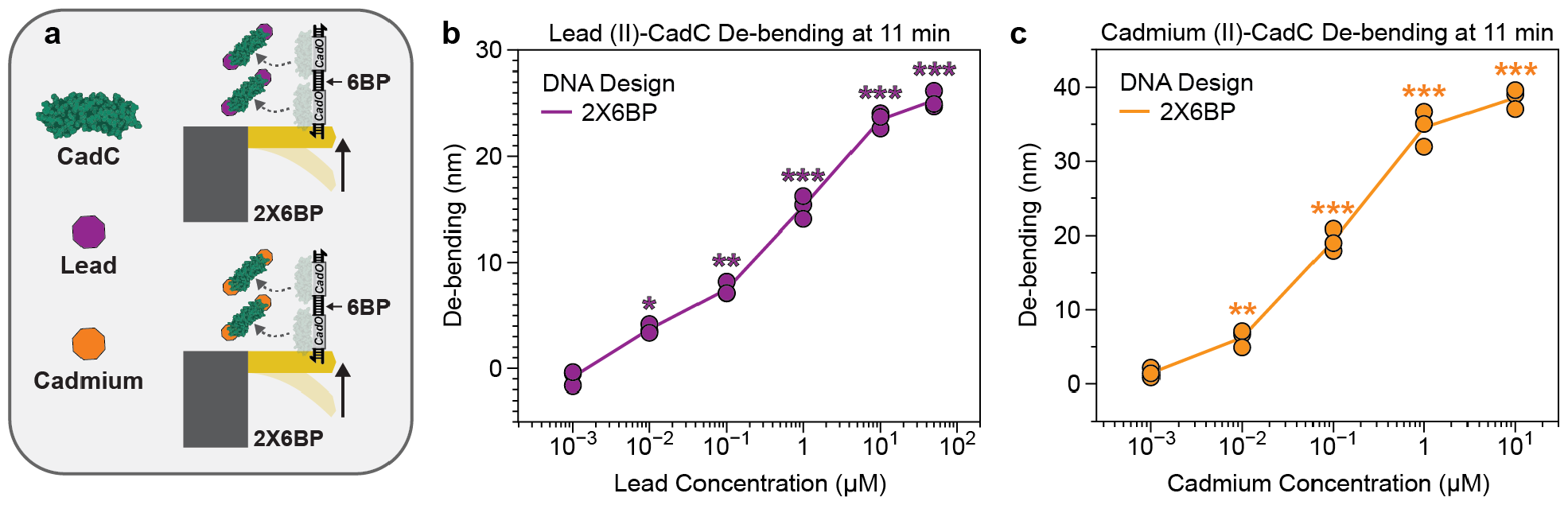
Microcantilevers interfaced with the CadC transcription factor can detect nanomolar concentrations of lead and cadmium. a) Schematic depicting the lead-CadC and cadmium-CadC sensing systems with the 2X6BP DNA design. b) Observed de-bending at 11 minutes for a range of PbCl_2_ concentrations. c) Observed de-bending at 11 minutes for a range of CdCl_2_ concentrations. De-bending is computed as the difference in bending between the sensing microcantilever, induced with lead or cadmium, and the mean of the control microcantilevers, induced with 1 μM carbenicillin as in **Fig. 1**. Data shown are for n = 3 technical replicates, where each technical replicate, corresponding to a single microcantilever, is represented by a point. Each solid line passes through the mean of the data points. The ligand concentrations at which the bending magnitude was distinguishable from the carbenicillin induction control were determined using a two-tailed, homoscedastic Student’s t-test against the carbenicillin control condition, and their *P* value ranges are indicated with asterisks for the 2X6BP case (\*\*\**P* < 0.001, \*\**P* = 0.001– 0.01, \**P* = 0.01–0.05). Time course data plots can be found in **Fig. S4, 5**, and full data and P values can be found in the **Supplementary Data**.

These results demonstrated that the aTF-microcantilever platform provides greater sensitivity for detection than the current versions of solution-based aTF detection.

## DISCUSSION

In this work, we successfully merged cell-free biosensing systems with nanomechanical detection systems to create a new platform for the rapid and sensitive detection of chemical compounds. By patterning microcantilevers with DNA containing the operator sites of allosteric transcription factors, we were able to achieve 10 nM limits of detection for tetracycline, lead, and cadmium within ∼11 minutes (**Fig. 2, 3**). This represents concentrations of 4 ppb for tetracycline, 2 ppb for lead, and 1 ppb for cadmium, which are approximately two orders of magnitude lower than what has been achievable with cell-free biosensing systems that use aTFs to control reporter gene expression as an output (*3*). In addition, the nanomechanical system is as fast as cell-free transcriptional biosensors that use DNA strand displacement readouts (*4*).

Importantly, our LOD of 2 ppb for lead aligns with the Illinois Department of Public Health’s lead testing requirement for standard analytical chemistry techniques, which states that the minimum detection limit of a chosen analytical testing method should not exceed 2 ppb (*30*). Our LOD for lead is also below the EPA action level of 15 ppb, which stipulates that water systems must take action via public education or lead service line replacement if 10% of the samples collected from households using that water source exceed this level (*25*). Our limit of detection for cadmium is below the EPA maximum contaminant level goal (MCLG) of 5 ppb for cadmium (*2*), which describes the level of a contaminant in drinking water below which there is no known or expected risk to health.

While the microcantilever system has superior sensing abilities over previously reported cell-free biosensing systems, there are a number of additional advantages and some potential disadvantages of the system. In terms of advantages, microcantilevers address another limitation of cell-free biosensing systems which is the lack of quantification. While genetic circuits can be added to cell-free biosensing systems to produce devices that yield semi-quantitative readouts (*4*), they do not produce digital readouts and thus cannot offer the same level of quantification that a microcantilever system coupled with on-board data processing can. In fact, if on-board data processing is used, microcantilever systems could be used to deliver results even faster by using short time signal transients to predict outputs signals. In addition, microcantilever devices can be easily multiplexed (*18*), potentially enabling several chemicals to be detected simultaneously. However, these advantages may come at the additional cost of manufacturing and deploying microcantilever systems, compared to the relative ease of manufacturing and distributing single-use cell-free biosensing systems (*10*).

While the microcantilever and cell-free biosensing systems utilize the same transcription factor-DNA operator pairs to detect chemicals, we did observe that they appear to have different sensing properties. Specifically, the dose-response curves obtained using the microcantilever platform did not contain a characteristic sharp inflection point that is typical of the dose-response curves obtained using the cell-free gene expression systems (*3*). To investigate this observation, we constructed a simple double-equilibrium model of the tetracycline-TetR system with the 1X DNA design (see **Supplementary Information**). We found that the model can reproduce results obtained via the ROSALIND cell-free setup (*3*) when the association constants available in the literature for the tetracycline-TetR system were used. However, reasonable agreement between the model predictions and the microcantilever dose-response curves required that the literature association constants between aTF and DNA be modified by several orders of magnitude, and the initial TetR concentration be lowered by two orders of magnitude. This seems to indicate that the physics of aTF-DNA binding/un-binding on a two-dimensional patterned surface is causing a significant perturbation to the biochemistry of aTF-DNA binding, though more work is needed to understand the mechanistic details of these differences.

We also note that this work is similar in spirit to recent work that couples cell-free biosensors with portable glucose meters for multiplexed, digital pathogen detection (*12*). To achieve digital readout from single-use cell-free biosensors, Amalfitano et al. utilized genetic circuits to transform the outputs of transcriptional biosensors into a glucose signal via glucogenic enzymes. The enzymatically generated glucose can then be read by low-cost glucose meters for digital sensing. This strategy would be broadly applicable to many transcription-based cell-free biosensing platforms, including those that leverage aTFs for analyte detection. In contrast to transcription-based cell-free biosensing platforms, the microcantilever-based sensing strategy outlined here directly converts aTF binding events into digital output, with the potential for continuous readout when the microcantilevers are coupled with MOSFET technology. The work outlined here thus employs a fundamentally different approach to signal transduction and reporting, providing a complementary solution to single-use biosensors that involve cell-free gene expression.

Overall, the merging of cell-free biosensors and microcantilever sensing represents a promising new platform that provides substantial value by enabling diagnostic systems that can address important public health challenges such as measuring water quality (*1*). The ability for MOSFET devices to implement rapid and sensitive detection of water contaminants is particularly promising, as these systems can be embedded in integrated circuits (*15*), potentially enabling their deployment in portable devices, and can be multiplexed (*17, 18*) to potentially allow the detection of multiple contaminants simultaneously. Furthermore, the microcantilever sensing framework should be readily translatable to sensing other chemicals for which aTFs are available (*31*). Incorporation of on-board data processing can further be used to process and communicate data on-the-fly, helping to make the system more robust (*32*) and enabling large-scale data-driven approaches to addressing public health challenges (*1*). With further development, we envision that this system can be combined with MOSFET technology (*15*) to pave the way for widespread incorporation of this platform within portable digital devices.

## MATERIALS AND METHODS

### Synthesis of allosteric transcription factors (aTFs)

Expression and purification of the TetR and CadC aTFs were performed as described in (*3*). Briefly, for TetR expression and purification, sequence-verified pET-28c plasmids were transformed into the Rosetta 2(DE3) pLysS *E. coli* strain for protein expression. Cultures were grown up and induced for overexpression with isopropyl β-d-1-thiogalactopyranoside. The cultures were then pelleted via centrifugation, resuspended in lysis buffer (10 mM Tris-HCl, pH 7.5–8.5, 500 mM NaCl, 1 mM tris(2-carboxyethyl)phosphine (TCEP) and protease inhibitor (cOmplete EDTA-free Protease Inhibitor Cocktail, Roche)), and then lysed via ultrasonication. Lysates were then centrifuged, and the supernatant was purified using His-tag affinity chromatography followed by size-exclusion chromatography. Collected proteins were concentrated and buffer exchanged (25 mM Tris-HCl, 100 mM NaCl, 1 mM TCEP, 50% glycerol, v/v) via centrifugal filtration. For CadC expression and purification, a pET-3d plasmid backbone was used. Purification was then done following precipitation with polyethyleneimine (PEI), precipitation with ammonium sulfate, cation-exchange chromatography, and size-exclusion chromatography similarly to what has been previously described (*27*). Protein concentrations were measured via the Qubit Protein Assay Kit, and protein purity and size were verified via SDS-PAGE gel. Purified proteins were stored at -20 °C. ATFs were diluted from stock to their indicated working concentrations with 18 MOhm water.

### DNA coating of microcantilevers

All DNA oligos (**Table S1**) were purchased from Integrated DNA Technologies (IDT), using either standard desalting or HPLC purification as indicated. DNA oligo stock solutions were made by dissolving DNA oligos from IDT in MilliQ water to yield a concentration of 500 μM. To generate working dsDNA solutions for microcantilever coating, equimolar ratios of 5’ C6 thiol modified DNA oligos (500 μM stock) and their complements (500 μM stock) were combined along with 5X annealing buffer (500 mM potassium acetate, 150 mM HEPES, pH 8.0) and diluted to the proper final volume using MilliQ water. The solutions were then annealed by heating to 95°C for 3 minutes inside a heat block and then allowing them to cool at room temperature for 1 hour. 5 μM dsDNA solution was chosen for microcantilever coating based on experimentation with variable concentrations of dsDNA (**Fig. S6**). Gold-coated microcantilevers purchased from Nanoworld Innovative Technologies (Product # ARROW-TL1Au-50), were then placed inside the wells of a 96 well plate containing the 5 μM dsDNA oligo solution (one microcantilever per well) and left to stand for approximately two hours at room temperature. The microcantilevers were then retrieved from the wells, air dried and used for analyte detection.

### Microfluidic chip design and manufacture

Microfluidic chip design and manufacture followed previously published protocols (*16, 33*). Microfluidic chips were designed and manufactured by erstwhile Nanoink Inc in 2006. These microfluidic chips were employed for detection purposes in our earlier published work (*16, 33*). Briefly, the microfluidic chips, essential components in microscale experiments, were intricately fabricated through processes such as photolithography and etching by Nanoink Inc in 2006.

### AFM deflection measurement for bending/de-bending reactions

For each bending reaction, a DNA-coated microcantilever was placed inside a fabricated microfluidic chip inside an atomic force microscope system (Bruker High Performance Bioscope Resolve Life Science Imaging System). The laser was aligned manually, and measurement initiated. 10 μL of allosteric transcription factor solution was then added to the microfluidic chip via pipette (see synthesis of allosteric transcription factors for details on aTF working solution preparation). Microcantilever deflection was then recorded at 1-minute intervals for a 15-minute period. For each de-bending reaction, a microcantilever was first bent using 10 μL of aTF solution (1 μM TetR or 5 μM CadC) as described in the procedure above. The 1 μM TetR and 5 μM CadC solutions used for de-bending studies were the same solutions used in the bending studies. Next, 10 μL of solution containing a specified ligand or control compound at the indicated concentration was added to the microfluidic chip. To control for the effect of the added ligand solution diluting the aTF concentration and potentially causing de-bending, we performed experiments with the non-cognate ligand carbenicillin which is not recognized by the aTFs used in this study. Microcantilever deflection was then recorded at 1-minute intervals for a 15-minute period.

### Calculation of microcantilever bending and de-bending

The measurement of microcantilever deflection was used to calculate bending and de-bending in different experimental conditions (**Fig. S2**). In the presence of aTF alone, bending was calculated as:

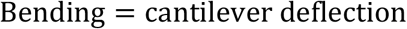

The calculation of de-bending combined deflection measurements in the presence of cognate aTF ligands and control compounds as follows (**Fig. S2**):

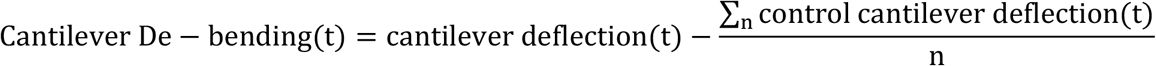

Where *cantilever deflection* is the measurement of deflection in the presence of cognate ligand, *control cantilever deflection* is the measurement of deflection in the presence of a control compound taken at the same time point as the cantilever deflection, and *n* is equal to the number of control cantilevers used.

The calculation of standard deviation (s) for use in error shading as evident in **Fig. 1b** was computed as follows for bending experiments:

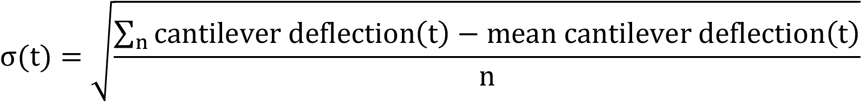

The calculation of standard deviation (σ_A−B_) for use in error shading as evident in **Fig. 1C** was computed as follows for de-bending experiments:

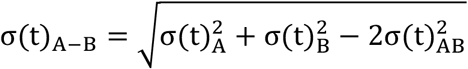

Where 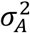is variance of deflection measurements at a given time in the presence of cognate ligand, 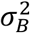is variance of deflection measurements in the presence of the control compound, and 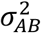 is their covariance.

## Supporting information

Jupyter Notebook

## STATISTICS AND REPRODUCIBILITY

The number of replicates and types of replicates performed are described in the legend to each figure. Individual data points are shown, and where relevant, the average ± standard deviation is shown; this information is provided in each figure legend. The type of statistical analysis performed in **Fig. 2c, 3b, and 3c** is described in the legend to each figure. Exact p-values computed from the statistical analysis can be found in the **Supplementary Data**.

## DATA AVAILABILITY

All data presented in this manuscript are available as supplementary data files.

## CODE AVAILABILITY

The Jupyter Notebook with the Python code used in Supplementary Note: Mechanistic Model of Cantilever-Based Detection Sensitivity Improvement, is provided as a supplementary file.

## AUTHOR CONTRIBUTIONS

D.K.A., J.K.J., G.S.S., J. B. L., V.P.D. designed the study. D.K.A, T.J.L, J. B. L. analyzed the data. D.K.A., J.K.J., G.S.S. conducted the research. D.K.A., J.K.J. developed the methodology. T.J.L undertook visualization of the data. D.K.A., T.J.L, J.K.J., J. B. L. wrote the article. D.K.A., T.J.L, J. B. L. curated the data. J. B. L., G.S.S., V.P.D. acquired funding for the study. D.K.A., G.S.S. validated the results. D.K.A., J. B. L. managed and coordinated the study. G.S.S., J. B. L., V.P.D. supervised the research.

## COMPETING INTERESTS STATEMENT

GSS, VPD and JBL have filed a provisional patent describing aspects of this work (application no. TBD). JBL is a founder, and has financial interests in, Stemloop Inc. The latter interests are reviewed and managed by Northwestern University in accordance with their conflicts of interest policies.

## ACKNOWLEDGEMENTS

This material is based upon work supported by a Northwestern McCormick Catalyst Award, and the National Science Foundation under Grant no. 2319427. In addition to this support, T.J.L. was supported in part by Northwestern University’s Synthesizing Biology Across Scales National Research Training Program (grant No. 2021900), and J.K.J. was supported by a Ryan Fellowship and by a Northwestern McCormick School of Engineering Terminal Year Fellowship. Any opinions, findings, and conclusions or recommendations expressed in this material are those of the authors and do not necessarily reflect the views of the National Science Foundation.

We thank Professor Daiana Capdevila and Matias Villarruel Dujovne, Fundación Instituto Leloir and Instituto de Investigaciones Bioquímicas de Buenos Aires (FIL-IIBBA, CONICET), Argentina, for providing the purified CadC protein used in this work.

## Notes

### Summary of Updates

Updated protein purification methods section and acknowledgements section.

## REFERENCES

1. C. J. Vorosmarty, P. B. McIntyre, M. O. Gessner, D. Dudgeon, A. Prusevich, P. Green, S. Glidden, S. E. Bunn, C. A. Sullivan, C. R. Liermann, P. M. Davies, Global threats to human water security and river biodiversity. Nature 467, 555–561 (2010).

2. (2023, 01/09/2023). “National Primary Drinking Water Regulations | US EPA.” Retrieved 12/05/2023, 2023, from https://www.epa.gov/ground-water-and-drinking-water/national-primary-drinking-water-regulations.

3. J. K. Jung, K. K. Alam, M. S. Verosloff, D. A. Capdevila, M. Desmau, P. R. Clauer, J. W. Lee, P. Q. Nguyen, P. A. Pasten, S. J. Matiasek, J. F. Gaillard, D. P. Giedroc, J. J. Collins, J. B. Lucks, Cell-free biosensors for rapid detection of water contaminants. Nat Biotechnol 38, 1451–1459 (2020).

4. J. K. Jung, C. M. Archuleta, K. K. Alam, J. B. Lucks, Programming cell-free biosensors with DNA strand displacement circuits. Nat Chem Biol 18, 385–393 (2022).

5. A. D. Silverman, U. Akova, K. K. Alam, M. C. Jewett, J. B. Lucks, Design and Optimization of a Cell-Free Atrazine Biosensor. ACS Synth Biol 9, 671–677 (2020).

6. W. Thavarajah, A. D. Silverman, M. S. Verosloff, N. Kelley-Loughnane, M. C. Jewett, J. B. Lucks, Point-of-Use Detection of Environmental Fluoride via a Cell-Free Riboswitch-Based Biosensor. ACS Synth Biol 9, 10–18 (2020).

7. T. Pellinen, T. Huovinen, M. Karp, A cell-free biosensor for the detection of transcriptional inducers using firefly luciferase as a reporter. Anal Biochem 330, 52–57 (2004).

8. K. Y. Wen, L. Cameron, J. Chappell, K. Jensen, D. J. Bell, R. Kelwick, M. Kopniczky, J. C. Davies, A. Filloux, P. S. Freemont, A Cell-Free Biosensor for Detecting Quorum Sensing Molecules in P. aeruginosa-Infected Respiratory Samples. ACS Synth Biol 6, 2293–2301 (2017).

9. M. P. McNerney, F. Piorino, C. L. Michel, M. P. Styczynski, Active Analyte Import Improves the Dynamic Range and Sensitivity of a Vitamin B(12) Biosensor. ACS Synth Biol 9, 402–411 (2020).

10. L. Zhang, W. Guo, Y. Lu, Advances in Cell-Free Biosensors: Principle, Mechanism, and Applications. Biotechnol J 15, e2000187 (2020).

11. W. Thavarajah, P. M. Owuor, D. R. Awuor, K. Kiprotich, R. Aggarwal, J. B. Lucks, S. L. Young, The accuracy and usability of point-of-use fluoride biosensors in rural Kenya. NPJ Clean Water 6, 5 (2023).

12. E. Amalfitano, M. Karlikow, M. Norouzi, K. Jaenes, S. Cicek, F. Masum, P. Sadat Mousavi, Y. Guo, L. Tang, A. Sydor, D. Ma, J. D. Pearson, D. Trcka, M. Pinette, A. Ambagala, S. Babiuk, B. Pickering, J. Wrana, R. Bremner, T. Mazzulli, D. Sinton, J. H. Brumell, A. A. Green, K. Pardee, A glucose meter interface for point-of-care gene circuit-based diagnostics. Nat Commun 12, 724 (2021).

13. G. Wu, R. H. Datar, K. M. Hansen, T. Thundat, R. J. Cote, A. Majumdar, Bioassay of prostate-specific antigen (PSA) using microcantilevers. Nat Biotechnol 19, 856–860 (2001).

14. A. K. Basu, A. Basu, S. Bhattacharya, Micro/Nano fabricated cantilever based biosensor platform: A review and recent progress. Enzyme Microb Technol 139, 109558 (2020).

15. G. Shekhawat, S. H. Tark, V. P. Dravid, MOSFET-Embedded microcantilevers for measuring deflection in biomolecular sensors. Science 311, 1592–1595 (2006).

16. D. K. Agarwal, A. C. Hunt, G. S. Shekhawat, L. Carter, S. Chan, K. Wu, L. Cao, D. Baker, R. Lorenzo-Redondo, E. A. Ozer, L. M. Simons, J. F. Hultquist, M. C. Jewett, V. P. Dravid, Rapid and Sensitive Detection of Antigen from SARS-CoV-2 Variants of Concern by a Multivalent Minibinder-Functionalized Nanomechanical Sensor. Anal Chem 94, 8105–8109 (2022).

17. Y. Min, H. Lin, D. E. Dedrick, S. Satyanarayana, A. Majumdar, A. S. Bedekar, J. W. Jenkins, S. Sundaram, A 2-D microcantilever array for multiplexed biomolecular analysis. Journal of Microelectromechanical Systems 13, 290–299 (2004).

18. G. S. Shekhawat, V. P. Dravid, Biosensors: Microcantilevers to lift biomolecules. Nat Nanotechnol 10, 830–831 (2015).

19. Code of Federal Regulations. (US General Services Administration, National Archives and Records Service, 2023).

20. C. A. Savran, S. M. Knudsen, A. D. Ellington, S. R. Manalis, Micromechanical detection of proteins using aptamer-based receptor molecules. Anal Chem 76, 3194–3198 (2004).

21. L. Cuthbertson, J. R. Nodwell, The TetR family of regulators. Microbiol Mol Biol Rev 77, 440–475 (2013).

22. K. Kotecka, A. Kawalek, M. Modrzejewska-Balcerek, J. Gawor, K. Zuchniewicz, R. Gromadka, A. A. Bartosik, Functional Characterization of TetR-like Transcriptional Regulator PA3973 from Pseudomonas aeruginosa. Int J Mol Sci 23, (2022).

23. P. Levallois, P. Barn, M. Valcke, D. Gauvin, T. Kosatsky, Public Health Consequences of Lead in Drinking Water. Curr Environ Health Rep 5, 255–262 (2018).

24. A. S. Winter, R. J. Sampson, From Lead Exposure in Early Childhood to Adolescent Health: A Chicago Birth Cohort. Am J Public Health 107, 1496–1501 (2017).

25. (09/01/2020). “Understanding the Lead and Copper Rule.” Retrieved 12/05/2023, 2023, from https://www.epa.gov/sites/default/files/2019-10/documents/lcr101_factsheet_10.9.19.final_.2.pdf

26. C. Rensing, Y. Sun, B. Mitra, B. P. Rosen, Pb(II)-translocating P-type ATPases. J Biol Chem 273, 32614–32617 (1998).

27. L. S. Busenlehner, N. J. Cosper, R. A. Scott, B. P. Rosen, M. D. Wong, D. P. Giedroc, Spectroscopic properties of the metalloregulatory Cd(II) and Pb(II) sites of S. aureus pI258 CadC. Biochemistry 40, 4426–4436 (2001).

28. G. Nucifora, L. Chu, T. K. Misra, S. Silver, Cadmium resistance from Staphylococcus aureus plasmid pI258 cadA gene results from a cadmium-efflux ATPase. Proc Natl Acad Sci U S A 86, 3544–3548 (1989).

29. G. Endo, S. Silver, CadC, the transcriptional regulatory protein of the cadmium resistance system of Staphylococcus aureus plasmid pI258. J Bacteriol 177, 4437–4441 (1995).

30. “Lead in Water: Background and Sampling Procedures.” Retrieved 12/05/2023, 2023, from https://dph.illinois.gov/content/dam/soi/en/web/idph/files/publications/publicationsdo-ohpidhp-liw-presentation.pdf.

31. R. Fernandez-Lopez, R. Ruiz, F. de la Cruz, G. Moncalian, Transcription factor-based biosensors enlightened by the analyte. Front Microbiol 6, 648 (2015).

32. S. Mostafa, I. Lee, S. K. Islam, S. A. Eliza, G. Shekhawat, V. P. Dravid, F. S. Tulip, Integrated MOSFET-Embedded-Cantilever-Based Biosensor Characteristic for Detection of Anthrax Simulant. IEEE Electron Device Letters 32, 408–410 (2011).

33. D. K. Agarwal, V. Nandwana, S. E. Henrich, V. Josyula, C. S. Thaxton, C. Qi, L. M. Simons, J. F. Hultquist, E. A. Ozer, G. S. Shekhawat, V. P. Dravid, Highly sensitive and ultra-rapid antigen-based detection of SARS-CoV-2 using nanomechanical sensor platform. Biosens Bioelectron 195, 113647 (2022).

