## Supplemental Information for "Ultra-sensitive water contaminant detection with transcription factor interfaced microcantilevers"

### Supplementary Information for Ultra-sensitive water contaminant detection with transcription factor interfaced microcantilevers

Dilip K. Agarwal[1], Tyler J. Lucci[2, 3, 4], Jaeyoung K. Jung[2, 3, 4,†], Gajendra S. Shekhawat[1], Julius B. Lucks [2, 3, 4, 5, 6,*] and Vinayak P. Dravid [1, 6, 7,*]

1 – Department of Material Science and Engineering and NU*ANCE* Center, Northwestern University, Evanston, IL 60208.

2 – Department of Chemical and Biological Engineering, Northwestern University (Evanston, IL, USA)

3 – Center for Synthetic Biology, Northwestern University (Evanston, IL, USA)

4 – Center for Water Research, Northwestern University (Evanston, IL, USA)

5 – Interdisciplinary Biological Sciences Graduate Program, Northwestern University (Evanston, IL, USA)

6 – Chemistry of Life Processes Institute, Northwestern University (Evanston, IL, USA)

7 – Robert H. Lurie Comprehensive Cancer Center, Northwestern University (Chicago, IL, USA)

[†]Present address: Mammoth Biosciences, Inc. (Brisbane, CA, USA)

**Mechanistic Model of Cantilever-Based Detection Sensitivity Improvement**

Model Setup:

The items below describe the double-equilibrium model developed for the tetracycline-TetR system with the 1X DNA design (**Fig. S7**) to investigate the differences between the dose-response curves obtained in this work and those obtained via the ROSALIND setup (*1*). The variables are assigned values according to literature values, experimental conditions, or best estimates as indicated.

Model assumptions:

- TetR exists only as a dimer.
- One tetracycline molecule binds to one TetR dimer.
- TetR dimer bound to tetracycline cannot bind DNA.
- One *tetO* site per DNA.

Equations:

The definitions of the equilibrium dissociation constants are as follows:

1) $K_{D,L} = \frac{[aTC][TetR]}{[aTC \cdot TetR]}$
2) $K_{D,1} = \frac{[DNA][TetR]}{[DNA \cdot TetR]}$

The equilibrium dissociation constant equations can then be incorporated into the species conservation equations for total DNA, aTC and TetR. Note, because the DNA is deposited on the cantilever surface, DNA concentrations are in moles per unit area, while moles per unit volume are used for TetR and aTC because those species can be released in the solution surrounding the cantilever.

3) $[DNA]_0 = [DNA] + [DNA \cdot TetR] = [DNA] + \frac{[DNA][TetR]}{K_{D,1}}$
4) $[aTC]_0 = [aTC] + [aTC \cdot TetR] = [aTC] + \frac{[aTC][TetR]}{K_{D,L}}$

5) $[\mathrm{TetR}]_0 V = [\mathrm{TetR}]V + [\mathrm{aTC}\cdot\mathrm{TetR}]V + [\mathrm{DNA}\cdot\mathrm{TetR}]A = [\mathrm{TetR}]V + \frac{[\mathrm{aTC}][\mathrm{TetR}]}{K_{D,L}}V + \frac{[\mathrm{DNA}][\mathrm{TetR}]}{K_{D,1}}A$

Solving the double equilibrium:

For a given experimental condition (specified by the input concentration of each molecular species), the fraction of DNA unbound by TetR is proportional to the output signal generated. Therefore, the goal of the double equilibrium analysis is to solve for $[\mathrm{DNA}]/[\mathrm{DNA}]_0$ as a function of $[\mathrm{TetR}]_0$ and $[\mathrm{aTC}]_0$. Rearranging equation (5), and writing [aTC] and [DNA] in terms of [TetR] using equations (4) and (3), respectively, yields:

6) $0 = [\mathrm{TetR}]V + \frac{[\mathrm{aTC}]_0[\mathrm{TetR}]}{K_{D,L}+[\mathrm{TetR}]}V + \frac{[\mathrm{DNA}]_0[\mathrm{TetR}]}{K_{D,1}+[\mathrm{TetR}]}A - [\mathrm{TetR}]_0 V$

Equation (6) can be solved numerically for [TetR], which can then be plugged into a rearranged version of equation (3) to find:

7) $\frac{[\mathrm{DNA}]}{[\mathrm{DNA}]_0} = \frac{1}{1+\frac{[\mathrm{TetR}]}{K_{D,1}}}$

For the ROSLAND system, all concentrations are on a volume basis, so equation (6) is re-written as:

8) $0 = [\mathrm{TetR}] + \frac{[\mathrm{aTC}]_0[\mathrm{TetR}]}{K_{D,L}+[\mathrm{TetR}]} + \frac{[\mathrm{DNA}]_0[\mathrm{TetR}]}{K_{D,1}+[\mathrm{TetR}]} - [\mathrm{TetR}]_0$

Making use of the fact that the volume term cancels. Solving (8) numerically can then be used in equation (7) in the same way.

Analysis of ROSALIND sensitivity:

We first explored the ability of the model to reproduce the results obtained using the ROSALIND setup (*1*). Specifically, we numerically solved equation (8) for the case of 25 nM DNA template and 1250 nM TetR, with other parameters specified as indicated in **Table S3**. Results from this model are in good agreement with those from experiment, with the model accurately predicting the aTC concentration at which the system activates (**Fig. S8**).

Analysis of microcantilever sensitivity:

We next used the double-equilibrium model to investigate the predicted dose-response curve for the microcantilever setup using parameter values from **Table S3**. Because the concentration of DNA immobilized on the microcantilever surface is unknown, we chose to evaluate two different DNA concentrations corresponding to ten times the maximum feasible concentration (F = 10) and one one-hundredth the maximum feasible concentration (F = 0.01). For the two DNA concentrations studied, the dose response curves predicted by the model are nearly identical. This is because in both cases, TetR is present in great excess compared to DNA, or $\frac{[\mathrm{TetR}]_0}{[\mathrm{DNA}]_0} \gg 1$, and because $\frac{K_{D,1}}{K_{D,L}} \gg 1$. The model's predictions are also of much different shape compared to the dose-response curves obtained via the microcantilever setup, although the EC50 values for the model and the data are generally within an order of magnitude of each other (**Fig. S9**).

We next explored if closer agreement between the model's dose-response output and de-bending data could be achieved by modifying the values of the variables in the model. Using the maximum feasible surface DNA concentration (F = 1), closer agreement between the model's dose-response output and de-bending data required lowering the initial concentration of TetR ($[\mathrm{TetR}]_0$) by approximately two orders of magnitude (from 500 nM to 5 nM) and lowering the TetR-*tetO* dissociation constant ($K_{D,1}$) by approximately four orders of magnitude (from 1.79e-10 to 2.27e-14 M) (**Fig. S10**). Lowering $K_{D,1}$ was required to achieve a shallower dose-response slope compared to the step-like response predicted using $K_{D,1}$ from literature, while lowering $[\mathrm{TetR}]_0$ was required to shift the EC50 value to better align with those of the de-bending data. While the concentration of aTF is indeed lowered during de-bending (10 μL of ligand solution is added to the existing 10 μL of aTF solution used for bending), this twofold decrease in aTF concentration is not nearly sufficient to account for the hundred-fold decrease in aTF concentration required for the model to better agree with the data.

The above results suggest that the microcantilever sensing system behaves differently from the ROSALIND sensing system (*1*), even though both systems employ the same aTF-based sensing strategy. One possible source of difference is signal propagation; in the case of the microcantilever, aTF binding and unbinding is transduced into surface stress and thus cantilever deflection, while for ROSALIND, aTF binding and unbinding is transduced into transcription and thus fluorescent aptamer formation.

It is also possible that aTF-DNA interactions are altered in the microcantilever setup, where DNA is immobilized on the microcantilever surface as opposed to diffusing freely in solution. Immobilizing DNA on the microcantilever surface could influence aTF-DNA interactions if the immobilized DNA forms a dense forest on the microcantilever surface that affects the ability of aTF to bind/unbind from the DNA, or if DNA structural flexibility is altered by its immobilization. The environment around the cantilever may also differ substantially from that of the bulk as it would be expected that the immobilized DNA would result in a highly localized negative charge that would need to be countered by the presence of other cations in the area, possibly resulting in a localized area of high ionic strength.

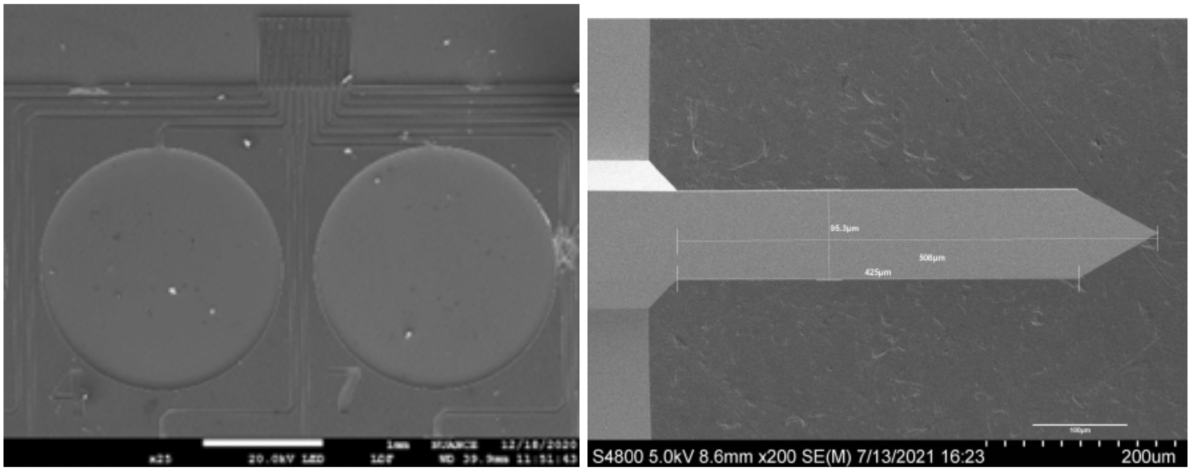


**Fig. S1. Microfluidic chip design and microcantilever.** An SEM image of the microfluidic chip design (left) and microcantilever (right) used for microcantilever bending and de-bending experiments based on previous work (*2, 3*). The microfluidic chips, essential components in microscale experiments, were intricately fabricated through processes such as photolithography and etching by Nanoink Inc in 2006.

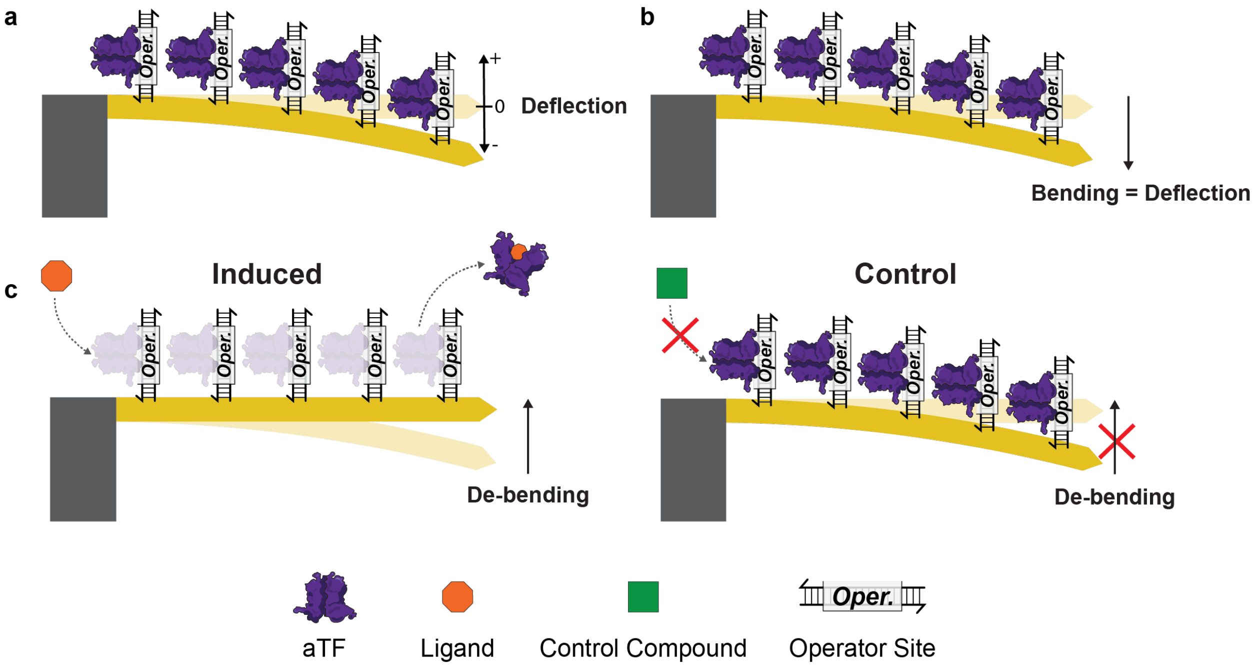


**Fig. S2. Schematic depiction of microcantilever bending and de-bending measurements.** A) Microcantilever deflection is measured as the distance of the tip of the microcantilever from the origin of the undeflected cantilever. B) aTF-DNA interactions cause microcantilever bending, where the distance the cantilever bends is equivalent to the measured deflection. C) Upon the addition of a ligand, aTF-ligand interactions cause an allosteric change, resulting in unbinding of the aTF and a de-bending of the cantilever (“induced,” left). The addition of a control compound that does not bind the aTF does not result in an allosteric change or microcantilever de-bending. The difference in microcantilever deflection between the ligand and control conditions is used to calculate ligand-induced microcantilever de-bending.

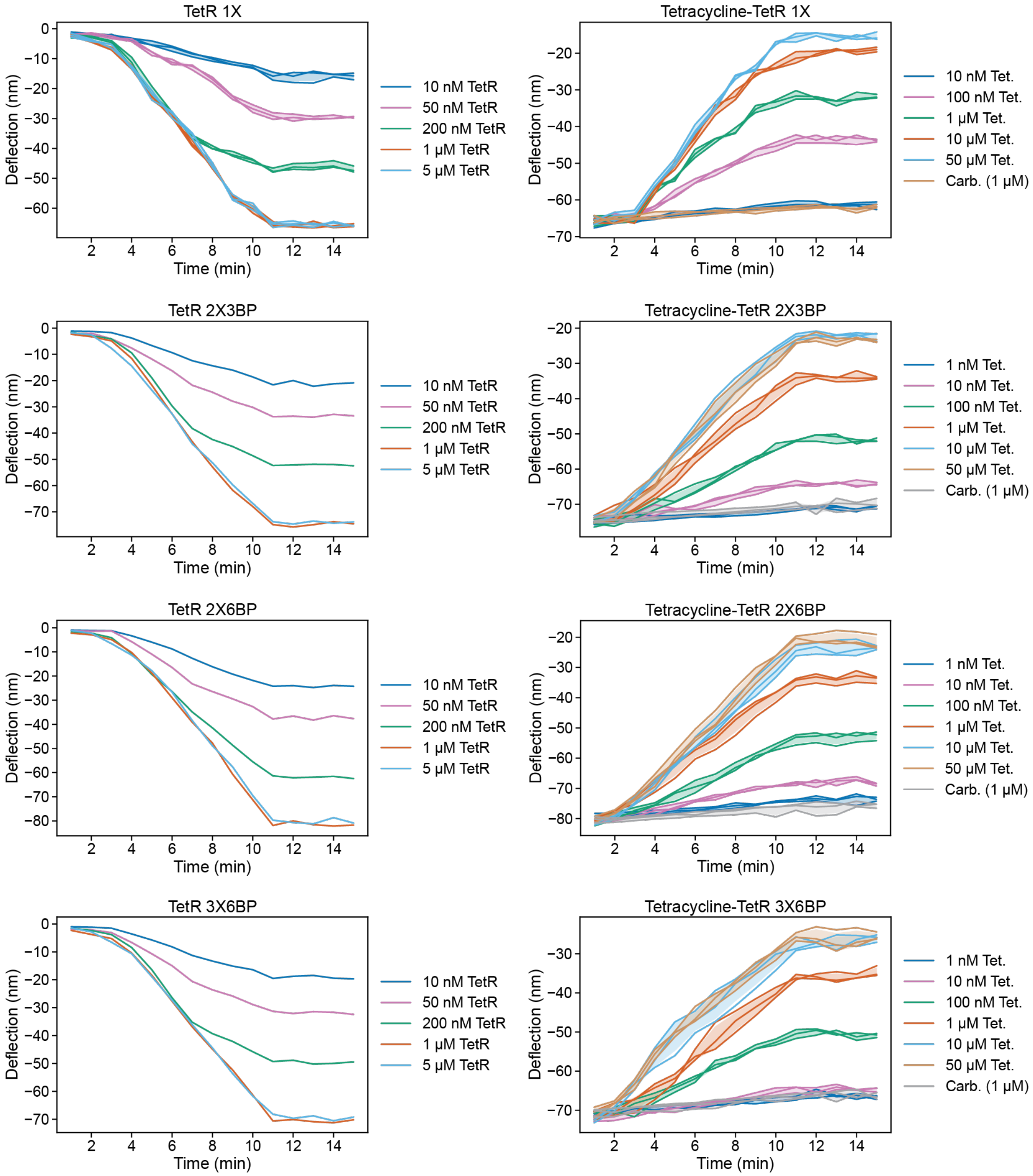


**Fig. S3. Raw AFM microcantilever deflection data for TetR bending and tetracycline-mediated de-bending using different combinations of *tetO* operator sites and spacers.** Left column: Bending in the presence of varying concentrations of TetR. Each plot is labeled with the DNA design used. For example, the DNA design "2X3BP" indicates two *tetO* operator sites per dsDNA with three base pair spacing between them, and the DNA design "1X" indicates one operator site for dsDNA. A list of all DNA designs used is in **Table S1**. Each solid line displayed in the plots corresponds to a single microcantilever's deflection over time as measured via AFM (one cantilever = one technical replicate), while the shaded region represents plus/minus one standard deviation about the mean of all technical replicates at each condition. For plots with no shaded region, only one cantilever was studied, and therefore standard deviation could not be computed. Right column: de-bending in the presence of either tetracycline ("Tet") or carbenicillin (“Carb). For de-bending experiments, 10 μL of 1 μM TetR was used in each experiment to first bend the cantilever, followed by addition of 10 μL of the indicated concentration of tetracycline or carbenicillin (see **Materials and Methods**). DNA design labels and plotting are the same as the left column.

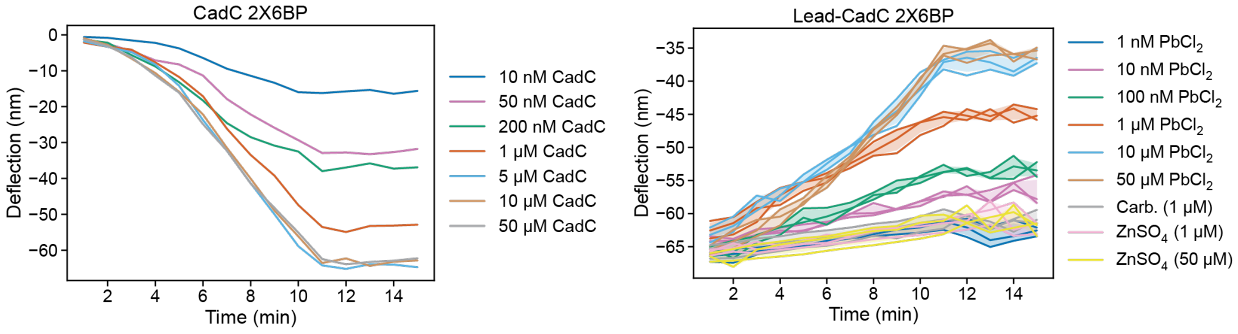


**Fig. S4. Raw microcantilever deflection data as measured by AFM for the sensor's response to CadC addition (left column) and addition of the indicated concentration of $PbCl_2$, $ZnSO_4$, or carbenicillin (after bending with 10 µL of 5 µM CadC, right column) for the 2X6BP DNA design.** We chose to test the sensor's response to Zn(II) here because CadC is known to respond to Zn(II), albeit to a lesser extent than to Pb(II) and Cd(II) (*4*). Each solid line displayed in the plots corresponds to a single microcantilever's bending or de-bending over time as measured via AFM (one cantilever = one technical replicate), while the shaded region represents plus/minus one standard deviation about the mean of all technical replicates at each condition. For plots with no shaded region, only one cantilever was studied, and therefore standard deviation could not be computed. In the plot legends, "Carb." Indicates carbenicillin.

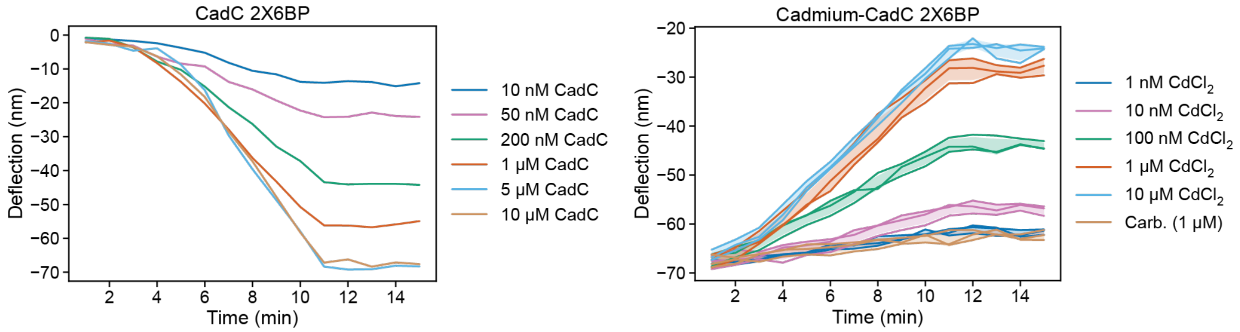


**Fig. S5**. **Raw microcantilever deflection data as measured by AFM for the sensor's response to CadC addition (left column) and addition of the indicated concentration of $CdCl_2$ or carbenicillin (after bending with 10 μL of 5 μM CadC, right column) for the 2X6BP DNA design.** Each solid line displayed in the plots corresponds to a single microcantilever's bending or de-bending over time as measured via AFM (one cantilever = one technical replicate), while the shaded region represents plus/minus one standard deviation about the mean of all technical replicates at each condition. For plots with no shaded region, only one cantilever was studied, and therefore standard deviation could not be computed. In the plot legends, "Carb." indicates carbenicillin.

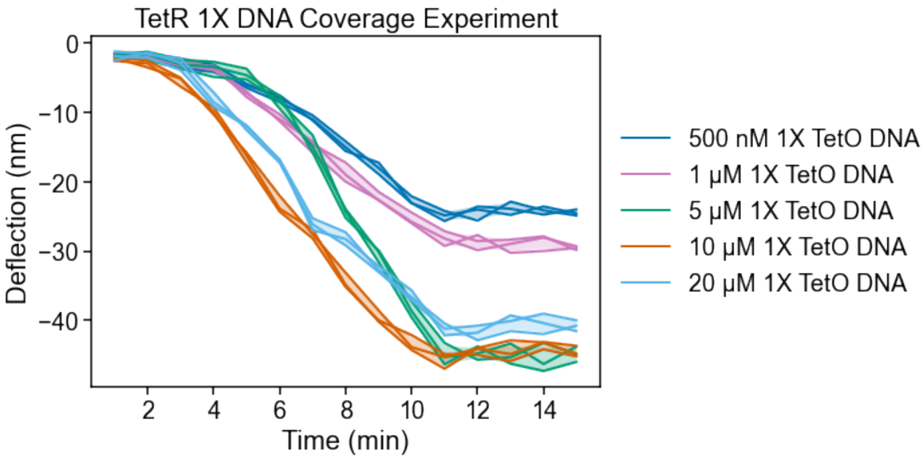


**Fig. S6**. **DNA coverage affects maximum realized deflection.** A plot displaying how microcantilever bending is affected by the concentration of DNA oligo used to coat the microcantilevers for the TetR system. The data above are plotted as measured by AFM using 10 μL of 1 μM TetR for bending and variable DNA oligo coating concentration. The DNA oligo used for this study is the "1X" *tetO* DNA oligo, meaning one TetR operator site per double-stranded DNA (dsDNA).

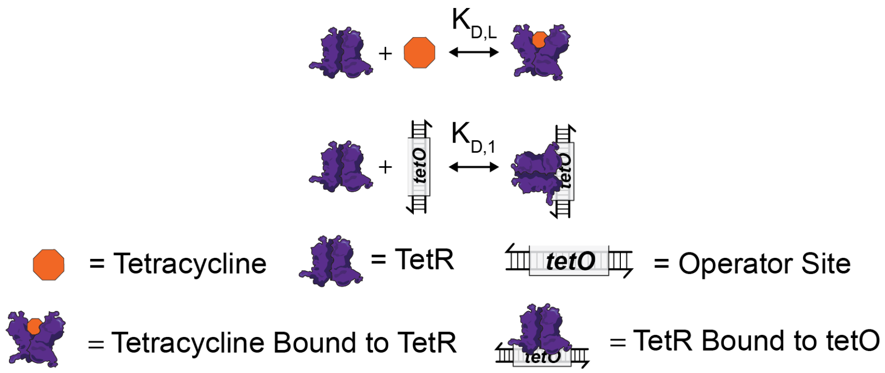


**Fig. S7. Double equilibrium model of transcription factor binding to DNA and target compound.** The double equilibrium model considers association/dissociation of tetracycline-TetR (top) and of TetR-*tetO* (bottom).

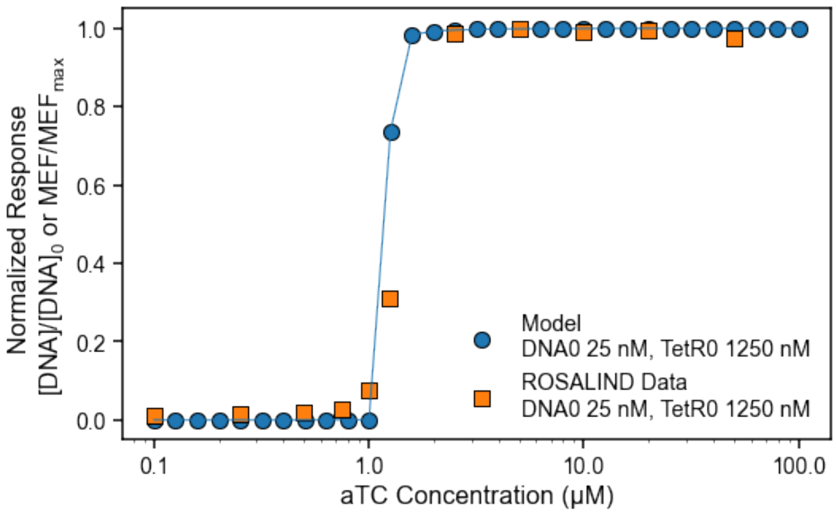


**Fig. S8. ROSALIND double equilibrium analysis.** Model prediction of fraction of DNA without TetR bound using the DNA template and TetR concentration studied in (*1*) with literature values for the association constants of tetracycline-TetR and TetR-*tetO*. The values used here are: $K_{D,L}$ = 7.94e-13 M, $K_{D,1}$ = 1.79e-10 M, $[DNA]_0 = 25$ nM, $[TetR]_0 = 1250$ nM, $[aTC]_0 = 0.1 - 100$ μM. Replotted induction curve (normalized) of the ROSALIND reaction from (*1*) using $[DNA]_0 = 25$ nM, $[TetR]_0 = 1250$ nM. Refer to (*1*) for methodological details of the ROSALIND experiment.

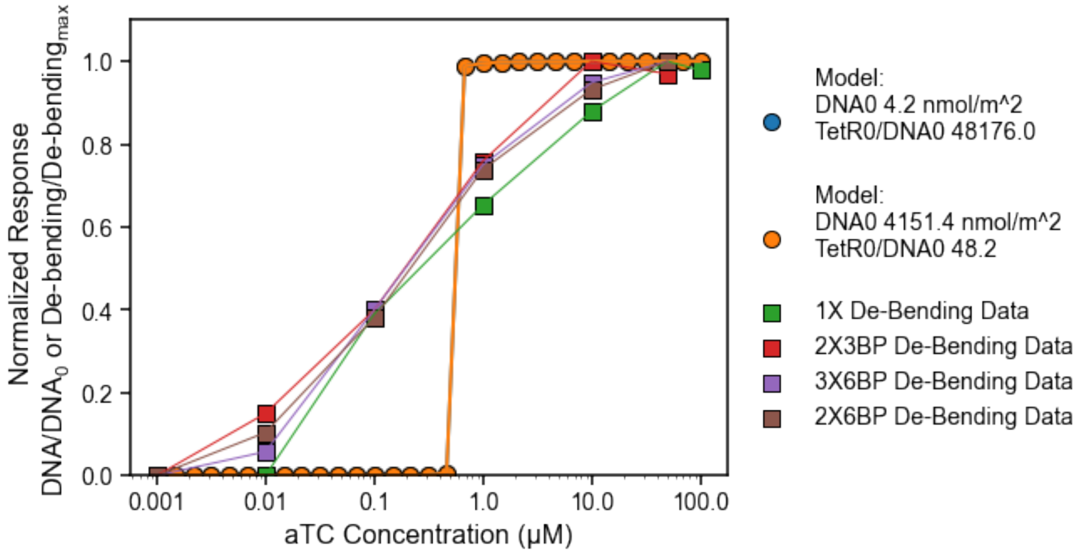


**Fig. S9. Base case microcantilever double equilibrium analysis.** A plot displaying the fraction of DNA without TetR bound as predicted by the model (circles) and an overlay of the normalized microcantilever de-bending data (squares) for the different DNA designs (see Fig. 2). The values used here are: $K_{D,L}$ = 7.94e-13 M, $K_{D,I}$ = 1.79e-10 M, $[\mathrm{DNA}]_0 = 4.2 - 4151\ \frac{\mathrm{nmol}}{\mathrm{m}^2}$, $[\mathrm{TetR}]_0 = 500\ \mathrm{nM}$, $[\mathrm{aTC}]_0 = 0.001 - 100\ \mu\mathrm{M}$. Two different values of $[\mathrm{DNA}]_0$ explored correspond to ten times the maximum feasible concentration (F = 10) and one one-hundredth the maximum feasible concentration (F = 0.01), where the maximum feasible concentration is based on the estimated area that a dsDNA molecule would occupy on the microcantilever surface. There is negligible difference in the model's prediction for the F = 10 and F = 0.01 cases and so those points overlap.

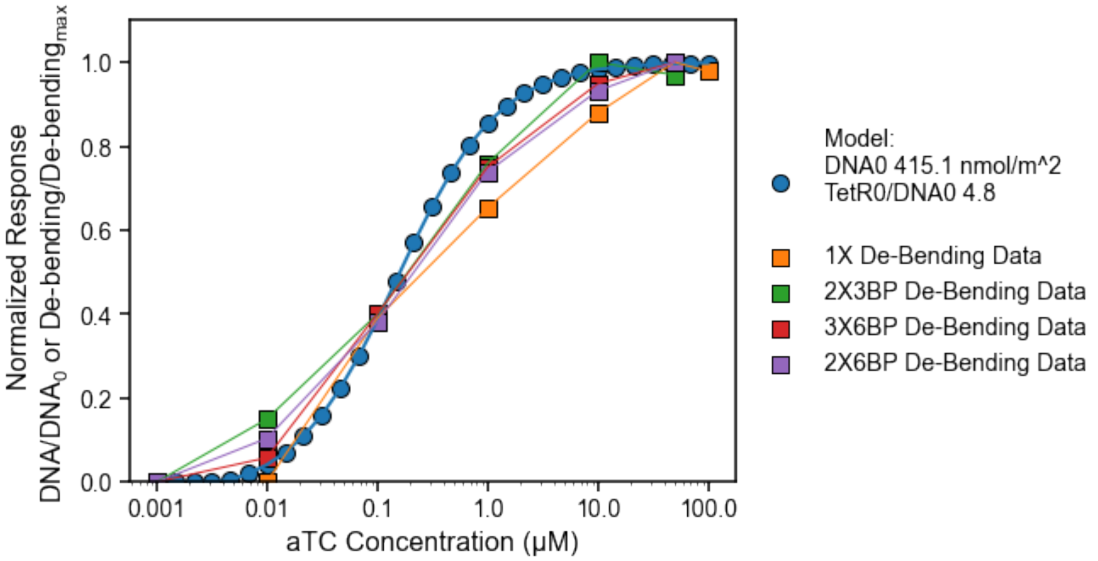


**Fig. S10. Changing TetR concentration and the TetR-DNA association constant generates closer agreement between the double equilibrium model and observed de-bending data.** A plot displaying the normalized microcantilever de-bending data (squares) for the different DNA designs along with an overlay of the fraction of DNA without TetR bound as predicted by the model (circles) using a lower initial TetR concentration (by two orders of magnitude) and a lower TetR-*tetO* dissociation constant (by ~four orders of magnitude). The values used here are: $K_{D,L}$ = 7.94e-13 M, $K_{D,1}$ = 2.27e-14 M, $[DNA]_0 = 415\ \frac{nmol}{m^2}$, $[TetR]_0 = 5$ nM, $[aTC]_0 = 0.001 - 100$ μM.

| | | |
|---|---|---|
| **Description** | 5' C6 thiol modified *tetO* sequence | |
| | **Sequence** | /5ThioMC6-D/GGA**TCCCTATCAGTGATAGAGA**CCC |
| | **Source** | IDT |
| | **Product** | 250 nmole DNA Oligo |
| | **Purification** | HPLC Purification |
| | **Used For** | TetR 1X DNA Design |
| **Description** | 5' C6 thiol modified *tetO* complement | |
| | **Sequence** | GGGTCTCTATCACTGATAGGGATCC |
| | **Source** | IDT |
| | **Product** | 25 nmole DNA Oligo |
| | **Purification** | Standard Desalting |
| | **Used For** | TetR 1X DNA Design |
| **Description** | 5' C6 thiol modified *tetO*X2 sequence | |
| | **Sequence** | /5ThioMC6-D/GGA**TCCCTATCAGTGATAGAGA****GGA****TCCCTATCAGTGATAGAGA**CCC |
| | **Source** | IDT |
| | **Product** | 250 nmole DNA Oligo |
| | **Purification** | HPLC Purification |
| | **Used For** | TetR 2X3BP DNA Design |
| **Description** | 5' C6 thiol modified *tetO*OX2 complement | |
| | **Sequence** | GGGTCTCTATCACTGATAGGGATCCTCTCTATCACTGATAGGGATCC |
| | **Source** | IDT |
| | **Product** | 25 nmole DNA Oligo |
| | **Purification** | Standard Desalting |
| | **Used For** | TetR 2X3BP DNA Design |
| **Description** | 5' C6 thiol modified *tetO*X2 sequence 6 BP spacer | |
| | **Sequence** | /5ThioMC6-D/ GGA**TCCCTATCAGTGATAGAGA****GGAGGA****TCCCTATCAGTGATAGAGA**CCC |
| | **Source** | IDT |
| | **Product** | 100 nmole DNA Oligo |
| | **Purification** | Standard Desalting |
| | **Used For** | TetR 2X6BP DNA Design |
| **Description** | 5' C6 thiol modified *tetO*X2 complement 6 BP spacer | |
| | **Sequence** | GGGTCTCTATCACTGATAGGGATCCTCCTCTCTATCACTGATAGGGATCC |
| | **Source** | IDT |
| | **Product** | 25 nmole DNA Oligo |
| | **Purification** | Standard Desalting |
| | **Used For** | TetR 2X6BP DNA Design |

| | | |
|---|---|---|
| **Description** | 5' C6 thiol modified *tetO*X3 sequence 6 BP spacer | |
| | **Sequence** | /5ThioMC6-D/ GGA**TCCCTATCAGTGATAGAGA****GGAGGA****TCCCTATCAGTGATAGAGA****GGAGGA****TCCCTATCAGTGATAGAGA**CCC |
| | **Source** | IDT |
| | **Product** | 100 nmole DNA Oligo |
| | **Purification** | Standard Desalting |
| | **Used For** | TetR 3X6BP DNA Design |
| **Description** | 5' C6 thiol modified *tetO*X3 complement 6 BP spacer | |
| | **Sequence** | GGGTCTCTATCACTGATAGGGATCCTCCTCTCTATCACTGATAGGGATCCTCCTCTCTATCACTGATAGGGATCC |
| | **Source** | IDT |
| | **Product** | 100 nmole DNA Oligo |
| | **Purification** | Standard Desalting |
| | **Used For** | TetR 3X6BP DNA Design |
| **Description** | 5' C6 thiol modified *cadO* sequence | |
| | **Sequence** | /5ThioMC6-D/ GGA**CTCAAATAAATATTTGAATGAA**CCC |
| | **Source** | IDT |
| | **Product** | 100 nmole DNA Oligo |
| | **Purification** | Standard Desalting |
| | **Used For** | CadC 1X DNA Design |
| **Description** | 5' C6 thiol modified *cadO* complement | |
| | **Sequence** | GGGTTCATTCAAATATTTATTTGAGTCC |
| | **Source** | IDT |
| | **Product** | 25 nmole DNA Oligo |
| | **Purification** | Standard Desalting |
| | **Used For** | CadC1X DNA Design |
| **Description** | 5' C6 thiol modified *cadO*X2 sequence 6BP spacer | |
| | **Sequence** | /5ThioMC6-D/ GGA**CTCAAATAAATATTTGAATGAA****GGAGGA****CTCAAATAAATATTTGAATGAA**CCC |
| | **Source** | IDT |
| | **Product** | 100 nmole DNA Oligo |
| | **Purification** | Standard Desalting |
| | **Used For** | CadC 2X6BP DNA Design |
| **Description** | 5' C6 thiol modified *cadO*X2 complement | |
| | **Sequence** | GGGTTCATTCAAATATTTATTTGAGTCCTCCTTCATTCAAATATTTATTTGAGTCC |
| | **Source** | IDT |
| | **Product** | 100 nmole DNA Oligo |
| | **Purification** | Standard Desalting |
| | **Used For** | CadC 2X6BP DNA Design |

**Table S1. DNA Oligos used in microcantilever experiments.** aTF operator sites are colored in **purple**, with base pair spacers highlighted in **orange**.

| Abbreviation | Description |
| --- | --- |
| $\mathrm{K_{D,L}}$ | Dissociation constant for tetracycline (aTC) and TetR |
| $\mathrm{K_{D,1}}$ | Dissociation constant for TetR and *tetO* site on DNA |
| $[\mathrm{DNA}]_0$ | Total dsDNA concentration on microcantilever surface (or in solution for ROSALIND) |
| $[\mathrm{aTC}]_0$ | Total tetracycline concentration (anhydrotetracycline) in sample chamber on microfluidic chip |
| $[\mathrm{TetR}]_0$ | Total TetR concentration in sample chamber on microfluidic chip |
| $\mathrm{A}$ | Area of microcantilever surface coated with DNA |
| $\mathrm{V}$ | Volume of liquid in sample chamber on microfluidic chip |
| $\mathrm{a}$ | Area occupied by a dsDNA immobilized on the microcantilever surface (used for estimating the maximum feasible DNA concentration on the microcantilever surface) |
| $\mathrm{F}$ | Dilution factor (if the microcantilever surface not completely saturated with DNA) |
| $\mathrm{N_A}$ | Avogadro's number |
| $[DNA]$ | Concentration of dsDNA without TetR bound |
| $[TetR]$ | Concentration of TetR not bound to tetracycline or DNA |
| $[aTC \cdot TetR]$ | Concentration of TetR bound to tetracycline |
| $[DNA \cdot TetR]$ | Concentration of TetR bound to DNA |

**Table S2. Notation used for the double-equilibrium model.**

| Parameter | Value | Units | Reference and Note |
| --- | --- | --- | --- |
| $K_{D,L}$ | 7.94e-13 | M | (*5*) **a** |
| $K_{D,1}$ | 1.79e-10 | M | (*5*) **b** |
| $A$ | 50000 | $\mu m^2$ | **c** |
| $V$ | 20 | μL | **d** |
| $a$ | 4 | $nm^2$ | (*6*) **e** |
| $F$ | Variable | Unitless | **f** |
| $N_A$ | 6.022e23 molecules/mole | Molecules/mole | |
| $[DNA]_0$ | $\frac{A}{\text{aTF size}} \cdot \frac{1}{N_A} \cdot \frac{1}{A} \cdot F$ | Moles/area | **g** |
| $[aTC]_0$ | Variable | M | |
| $[TetR]_0$ | Variable | M | |

**Table S3. Parameters used in the double-equilibrium model.**

Notes:

a. Taken as the reciprocal of association constant for aTC and TetR, which is stated in (*5*)
b. Taken as the reciprocal of association constant for TetR and *tetO* site on DNA, which is stated in (*5*)
c. The gold-coated microcantilever surface area was calculated using an SEM image of a microcantilever.
d. The final (solution) volume of the microcantilever system is 20 μL (10 μL of aTF solution + 10 μL ligand solution).
e. The area occupied by a DNA molecule immobilized on the microcantilever surface was taken to be the square of the diameter of a short dsDNA fragment. The diameter of a short dsDNA fragment can be found in (*6*).
f. Because the actual DNA concentration on the microcantilever surface is unknown, a scaling factor, F, is used to explore different concentrations of DNA on the microcantilever surface. The scaling factor multiplies the maximum feasible DNA concentration on the microcantilever surface, which is computed as the maximum possible concentration of dsDNA that could be fit onto the microcantilever surface (microcantilever surface area divided by dsDNA area, a).
g. The total concentration of DNA on the microcantilever surface is computed by multiplying the maximum feasible DNA concentration by the scaling factor, F.

**REFERENCES**

1. J. K. Jung, K. K. Alam, M. S. Verosloff, D. A. Capdevila, M. Desmau, P. R. Clauer, J. W. Lee, P. Q. Nguyen, P. A. Pasten, S. J. Matiasek, J. F. Gaillard, D. P. Giedroc, J. J. Collins, J. B. Lucks, Cell-free biosensors for rapid detection of water contaminants. *Nat Biotechnol* **38**, 1451-1459 (2020).
2. D. K. Agarwal, A. C. Hunt, G. S. Shekhawat, L. Carter, S. Chan, K. Wu, L. Cao, D. Baker, R. Lorenzo-Redondo, E. A. Ozer, L. M. Simons, J. F. Hultquist, M. C. Jewett, V. P. Dravid, Rapid and Sensitive Detection of Antigen from SARS-CoV-2 Variants of Concern by a Multivalent Minibinder-Functionalized Nanomechanical Sensor. *Anal Chem* **94**, 8105-8109 (2022).
3. D. K. Agarwal, V. Nandwana, S. E. Henrich, V. Josyula, C. S. Thaxton, C. Qi, L. M. Simons, J. F. Hultquist, E. A. Ozer, G. S. Shekhawat, V. P. Dravid, Highly sensitive and ultra-rapid antigen-based detection of SARS-CoV-2 using nanomechanical sensor platform. *Biosens Bioelectron* **195**, 113647 (2022).
4. C. Rensing, Y. Sun, B. Mitra, B. P. Rosen, Pb(II)-translocating P-type ATPases. *J Biol Chem* **273**, 32614-32617 (1998).
5. A. Kamionka, J. Bogdanska-Urbaniak, O. Scholz, W. Hillen, Two mutations in the tetracycline repressor change the inducer anhydrotetracycline to a corepressor. *Nucleic Acids Res* **32**, 842-847 (2004).
6. H. Lederer, R. P. May, J. K. Kjems, G. Baer, H. Heumann, Solution structure of a short DNA fragment studied by neutron scattering. *Eur J Biochem* **161**, 191-196 (1986).
