## Supplementary material for "Ultra-sensitive water contaminant detection with transcription factor interfaced microcantilevers": Jupyter Notebook: Agarwal_Cantilever_Biosensing_of_Water_Contaminants_Jupyter_Notebook.ipynb

```
# %% [markdown]
## Supplementary Information for Rapid, ultra-sensitive detection of water contaminants with allosteric transcription factor interfaced microcantilevers

Dilip K. Agarwal$^{1}$, Tyler J. Lucci$^{2,3,4}$, Jaeyoung K. Jung$^{2,3,4}$, Gajendra S. Shekhawat$^{1}$, Julius B. Lucks$^{2,3,4,5,6,*}$, and Vinayak P. Dravid$^{1,6,7,*}$

1 – Department of Material Science and Engineering and NUANCE Center, Northwestern University, Evanston, IL 60208<br>
2 – Department of Chemical and Biological Engineering, Northwestern University, Evanston, IL, USA.<br>
3 – Center for Synthetic Biology, Northwestern University, Evanston, IL, USA.<br> 
4 – Center for Water Research, Northwestern University, Evanston, IL, USA<br>
5 – Interdisciplinary Biological Sciences Graduate Program, Northwestern University, Evanston, IL, USA<br> 
6 – Chemistry of Life Processes Institute, Northwestern University, Evanston, IL, 60208, USA<br> 
7 – Robert H. Lurie Comprehensive Cancer Center, Northwestern University, Chicago, IL, 60611, USA<br> 
*Corresponding author: Julius B. Lucks and Vinayak P. Dravid<br> 

# %% [markdown]
## Analysis of ROSALIND Sensitivity
The equation below is used to solve for [TetR] in the double equilibrium model for predicting the dose-response of the tetracycline-TetR system for the ROSALIND setup. Remaining variables are then solved for via back substitution. **See Supplementary Note for the full model and equations** <br> 
$0 = [TetR] + \frac{{[aTC]_0}[TetR]}{K_{D,L}+[TetR]} + \frac{{[DNA]_0}[TetR]}{K_{D,1}+[TetR]} - {[TetR]_0}$ <br>

# %% [code]
import numpy as np
import pandas as pd
import xlrd
import matplotlib
import matplotlib.pyplot as plt
import matplotlib.font_manager as font_manager
from scipy.optimize import root_scalar

# %% [code]
#returns x value that makes function 0 by linear interpolation
def interpolate_zero(x_vector,y_vector):
    val_zero = 0
    for k in range(1,len(x_vector)-1):
        y_ = y_vector[k]
        y_prev = y_vector[k-1]
        x_ = x_vector[k]
        x_prev = x_vector[k-1]
        #if function value is zero at this x, return the value of that x
        if y_ == 0:
            return x_
        #if function value is positive but previous value was negative, interpolate 
        elif y_ >= 0 and y_prev < 0:
            val_zero = x_prev+(0-y_prev)*(x_-x_prev)/(y_-y_prev)
            return val_zero
        #if function value is negative but previous value was positive, interpolate 
        elif y_ <= 0 and y_prev > 0:
            val_zero = x_prev+(0-y_prev)*(x_-x_prev)/(y_-y_prev)
            return val_zero
    return 'val_zero'

# %% [code]
#double-equilibrium function to be zeroed for 3D, 1X DNA case 
def three_D_one_x_DNA(x, bb, cc, dd, ee, ff):
    #[TetR] = x
    #[aTC]0 = bb
    #KDL = cc
    #[DNA]0 = dd
    #KD1 = ee
    #[TetR]0 = ff
    return x+bb*x/(cc+x)+dd*x/(ee+x) - ff

# %% [code]
#Function importing ROSALIND data, which can be found associated with the publication Cell-free biosensors for rapid detection of water contaminants by Jung et al
#This function returns the tetracycling-TetR data found in Figure 2h of Jung et al (averages plotted)
#Note: THE FILE NEEDS TO BE SAVED IN .XLS FORMAT (instead of .xlsx)
def import_three_D_one_x_DNA_ROSALIND(ROS_file_path):
    book = xlrd.open_workbook(ROS_file_path)
    nsheets = book.nsheets
    sheetNames = book.sheet_names()
    sheet = book.sheet_by_name('Fig2h')
    nrows = sheet.nrows
    ncols = sheet.ncols
    #array containing all values in the last row of the spreadsheet (which was plotted in Figure 2h)
    all_data_values = []
    #array containing the tetracycline concentrations (µM) corresponding to each condition
    tetracycline_array = []
    for col in range(1,ncols):
        all_data_values.append(sheet.cell(nrows-1,col).value)
        tetracycline_condition = sheet.cell(2,col).value
        if tetracycline_condition != '':
            splits = tetracycline_condition.split(' ')
            tetracycline_uM = float(splits[0])
            tetracycline_array.append(tetracycline_uM)
    n_replicates = 3
    #each condition has three replicates, so average values for each condition and make a new array
    average_data_values = []
    num_conditions = int(len(all_data_values)/n_replicates)
    for i in range(0,num_conditions):
        vals = all_data_values[i*3:i*3+3]
        average = np.mean(vals)
        average_data_values.append(average)
    #normalize data
    max_data_value = np.max(average_data_values)
    average_data_values_normalized = [yi/max_data_value for yi in average_data_values]
    return [tetracycline_array,average_data_values_normalized]

# %% [code]
KDL = 1/(1.26e12)*10**9; #nM PMID 14764926
KD1 = 1/(5.6e9)*10**9; #nM PMID 14764926

#TetR concentration range
TetR0 = 1250 #nM

#aTC0 (anhydrous tetracycline) concentration range
apoints = 31
aTC0_vector = np.logspace(2,5,apoints) #nM, 2 and 5 for ROSALIND

#DNA concentration range
DNA0 = 25; #nM

#convert aTC0 from nM to uM for plotting purposes
aTC0_for_plot = []
for i in range(0,len(aTC0_vector)):
    aTC0_for_plot.append(aTC0_vector[i]/1000)

#loop over all KDL/KD0 ratios
fig, ax = plt.subplots(figsize = (8,5))
fig.patch.set_facecolor('white')
fig.patch.set_alpha(0)
ax.patch.set_facecolor('white')
ax.patch.set_alpha(0)

#solve nonlinear equation for [aTF], backsolve for all other variables 
vals = []
for j in range(0,len(aTC0_vector)):
    aTC0 = aTC0_vector[j]
    #compute TetR concentration via linear interpolation to use as an initial guess to feed to Scipy root_scalar
    xvals = np.logspace(-21,np.log10(TetR0)+1,1001)
    yvals = []
    for i in range(0,len(xvals)):
        TetR = xvals[i]
        function_value = three_D_one_x_DNA(TetR, aTC0, KDL, DNA0, KD1, TetR0)
        yvals.append(function_value)
    initial_guess = interpolate_zero(xvals,yvals)
    #compute TetR concentration via Scipy root_scalar
    sol = root_scalar(three_D_one_x_DNA, args=(aTC0, KDL, DNA0, KD1, TetR0), bracket=[0, TetR0],
                     x0 = initial_guess)
    #compare which root is better (i.e. closer to 0)
    TetR = initial_guess
    func_value_for_initial_guess = three_D_one_x_DNA(initial_guess, aTC0, KDL, DNA0, KD1, TetR0)
    func_value_for_root_scalar_root = three_D_one_x_DNA(sol.root, aTC0, KDL, DNA0, KD1, TetR0)
    if func_value_for_root_scalar_root**2 < func_value_for_initial_guess**2:
        TetR = sol.root
    #backsolve for all other species concentrations
    aTC = aTC0/(1+TetR/KDL)
    aTCTetR = aTC*TetR/KDL
    DNA = DNA0/(1+TetR/KD1)
    DNATetR = TetR*DNA/KD1
    vals.append(DNA/DNA0)
    
#plot dose-response curve for specific DNA concentration
label = f'Model\nDNA0 {str(DNA0)} nM, TetR0 {str(TetR0)} nM'
ax.scatter(aTC0_for_plot, vals, s=100, marker='o', edgecolor='black', linewidth = 1, label = label)
#get normalized ROSALIND data for 25 nM DNA0, 1250 nM TetR0 and plot
fp = r'/Users/tjl7324/Documents/PhD/Projects/Cantilever/41587_2020_571_MOESM10_ESM.xls'
ROSALIND_data = import_three_D_one_x_DNA_ROSALIND(fp)
ax.scatter(ROSALIND_data[0], ROSALIND_data[1], s=100, marker='s', edgecolor='black', linewidth = 1,
           label = 'ROSALIND Data\nDNA0 25 nM, TetR0 1250 nM')
axes_font_size = 16
title_font_size = 16
legend_font_size = 14
font_name = 'Arial'
font_weight = 'normal'
ax.set_xlabel('aTC Concentration (\u03BCM)',fontsize = axes_font_size,fontweight = font_weight, fontname = font_name)
ax.set_ylabel('Normalized Response \n$\mathregular{[DNA]/[DNA]_{0}}$ or $\mathregular{MEF/MEF_{max}}$',fontsize = axes_font_size,fontweight = font_weight, fontname = font_name)
legend_font = font_manager.FontProperties(family=font_name,
                       weight=font_weight,
                       style='normal', 
                        size=legend_font_size)
ticks_font = font_manager.FontProperties(family=font_name,
                   weight=font_weight,
                   style='normal', 
                    size=legend_font_size)
tkw = dict(size = 6, width = 1.5)
ax.tick_params(axis ='y',
         **tkw)
ax.tick_params(axis ='x',
         **tkw)
ax.plot(aTC0_for_plot, vals, linewidth = 1)
ax.set_xscale('log')
#ax.legend(loc="lower right",prop={'size': 12})
leg = ax.legend(frameon=False,prop=legend_font,loc='lower right')
for label in ax.get_xticklabels():
    label.set_fontproperties(ticks_font)
for label in ax.get_yticklabels():
    label.set_fontproperties(ticks_font)
for line in leg.get_lines():
    line.set_linewidth(line_width_for_plot)
for axis in ['top', 'bottom', 'left', 'right']:
    ax.spines[axis].set_linewidth(1.5)  # change width
    ax.spines[axis].set_color('black')
ax.get_xaxis().set_major_formatter(matplotlib.ticker.FuncFormatter(lambda x, p: format(np.around(x,3), ',')))
plt.show()
    

# %% [markdown]
## Analysis of Microcantilever Sensitivity
The equation below is used to solve for [TetR] in the double equilibrium model for predicting the dose-response of the tetracycline-TetR system for the microcantilever setup. Remaining variables are then solved for via back substitution. **See Supplementary Note for the full model and equations** <br> 
$0 = [TetR]V + \frac{{[aTC]_0}[TetR]}{K_{D,L}+[TetR]}V + \frac{{[DNA]_0}[TetR]}{K_{D,1}+[TetR]}A - {[TetR]_0}V$ <br>

# %% [code]
#double-equilibrium function to be zeroed for 3D, 1X DNA case 
def two_D_variable_x_DNA(x, bb, cc, dd, ee, ff, vv, aa):
    #[TetR] = x
    #[aTC]0 = bb
    #KDL = cc
    #[DNA]0 = dd
    #KD1 = ee
    #[TetR]0 = ff
    return x*vv+bb*x/(cc+x)*vv+dd*x/(ee+x)*aa - ff*vv

# %% [code]
#data for 1X tetracycline cantilever, see Supplementary Data 1, spreadsheet tab "Tetracycline-1X-Debend"
#all values are at t = 11 min
sample_data_uM_tet_1X = [0.01,0.1,1,10,50,100]
sample_data_de_bending_1X = [1.06,18.74,30.70,40.97,46.46,45.49]

#data for 2X3BP tetracycline cantilever, see Supplementary Data 1, spreadsheet tab "Tetracycline-2X3BP-Debend"
#all values are at t = 11 min
sample_data_uM_tet_2X3BP = [0.001,0.01,0.1,1.0,10,50]
sample_data_de_bending_2X3BP = [-1.02,6.33,18.55,36.19,48.00,46.51]

#data for 3X6BP tetracycline cantilever, see Supplementary Data 1, spreadsheet tab "Tetracycline-3X6BP-Debend" 
#all values are at t = 11 min
sample_data_uM_tet_3X6BP = [0.001,0.01,0.1,1.0,10,50]
sample_data_de_bending_3X6BP = [-0.45,1.93,16.04,30.48,38.68,40.80]

#data for 2X6BP tetracycline cantilever, see Supplementary Data 1, spreadsheet tab "Tetracycline-2X6BP-Debend" 
#all values are at t = 11 min
sample_data_uM_tet_2X6BP = [0.001,0.01,0.1,1,10,50]
sample_data_de_bending_2X6BP = [2.17,7.69,22.40,41.41,51.67,55.29]

#data labels
data_labels = ['1X De-Bending Data','2X3BP De-Bending Data','3X6BP De-Bending Data','2X6BP De-Bending Data']

#normalize data
cantilever_x_datas = [sample_data_uM_tet_1X,sample_data_uM_tet_2X3BP,sample_data_uM_tet_3X6BP,sample_data_uM_tet_2X6BP]
cantilever_y_datas = [sample_data_de_bending_1X,sample_data_de_bending_2X3BP,sample_data_de_bending_3X6BP,sample_data_de_bending_2X6BP]
cantilever_normalized_y_datas = []
for i in range(0,len(cantilever_y_datas)):
    y_data = cantilever_y_datas[i]
    maxy = max(y_data)
    miny = min(y_data)
    normalized_y_data = []
    for j in range(0,len(y_data)):
        normalized_y_data.append((y_data[j]-miny)/(maxy-miny))
    cantilever_normalized_y_datas.append(normalized_y_data)

#define parameters for the model
KDL = 1/(1.26e12); #M PMID 14764926
KD1 = 1/(5.6e9); #M PMID 14764926
V = 20e-6 #L volume in cantilever chamber
A = 50000e-12 #m^2 area of cantilever
DNA_size = 2e-9 #m diameter of a dsDNA
NA = 6.022e23 #avagadro number molec/mole
F_vector = np.logspace(-2,1,2) #dilution factor for surface concentration of DNA
DNA0_max = A/DNA_size**2/NA/A #mol/m^2 maximum theoretical DNA concentration on microcantilever surface
print(f'max DNA: {np.around(DNA0_max*10**9,3)} nmol/m^2')
TetR0 = 500e-9; #M concentration of TetR in the system

#define aTC0 concentration range for computing dose-response
apoints = 31;
aTC0_vector = np.logspace(-9,-4,apoints); #M

#convert units of aTC0 for plotting purposes
aTC0_for_plot = []
for i in range(0,len(aTC0_vector)):
    aTC0_for_plot.append(aTC0_vector[i]*10**6)

#initialize figure
fig, ax = plt.subplots(figsize = (8,5))

#uncomment the line below to explore case where KD10 and [TetR]0 are changed to improve model/data agreement
#KD1 = 2.27e-14 #M
#F_vector = [1]
#TetR0 = 5e-9

#solve nonlinear equation for [aTF], backsolve for all other variables 
for nu in range(0,len(F_vector)):
    DNA0 = DNA0_max*F_vector[nu]
    print(DNA0)
    TetR_DNA_molar_ratio = (TetR0*V)/(DNA0*A)
    vals = [];
    #loop over all aTC0 concentrations and compute value of function to be zeroed
    for j in range(0,len(aTC0_vector)):
        aTC0 = aTC0_vector[j]
        #compute TetR concentration via linear interpolation to use as an initial guess to feed to Scipy root_scalar
        xvals = np.logspace(-21,np.log10(TetR0)+1,1001)
        yvals = []
        for i in range(0,len(xvals)):
            TetR = xvals[i]
            function_value = two_D_variable_x_DNA(TetR,aTC0, KDL, DNA0, KD1, TetR0, V, A)
            yvals.append(function_value)
        initial_guess = interpolate_zero(xvals,yvals)
        #compute TetR concentration via Scipy root_scalar
        sol = root_scalar(two_D_variable_x_DNA, args=(aTC0, KDL, DNA0, KD1, TetR0, V, A), bracket=[0, TetR0],
                         x0 = initial_guess)
        #compare which root is better (i.e. closer to 0)
        TetR = initial_guess
        func_value_for_initial_guess = two_D_variable_x_DNA(initial_guess, aTC0, KDL, DNA0, KD1, TetR0, V, A)
        func_value_for_root_scalar_root = two_D_variable_x_DNA(sol.root, aTC0, KDL, DNA0, KD1, TetR0, V, A)
        if func_value_for_root_scalar_root**2 < func_value_for_initial_guess**2:
            TetR = sol.root
        #backsolve for all other species concentrations
        aTC = aTC0/(1+TetR/KDL)
        aTCTetR = aTC*TetR/KDL
        DNA = DNA0/(1+TetR/KD1)
        DNATetR = TetR*DNA/KD1
        vals.append(DNA/DNA0)
    #plot dose-response curve for specific DNA concentration
    DNA_label = np.around(DNA0*10**9,1)
    TetR_DNA_molar_ratio_label = np.around(TetR_DNA_molar_ratio,1)
    label = f'Model: \nDNA0 {str(DNA_label)} nmol/m^2 \nTetR0/DNA0 {TetR_DNA_molar_ratio_label} \n'
    ax.scatter(aTC0_for_plot, vals, s=100, marker='o', edgecolor='black', linewidth = 1, label = label)
    ax.plot(aTC0_for_plot, vals, linewidth = 2)
    
#plot normalized microcantilever de-bending data
for k in range(0,len(cantilever_x_datas)):
    ax.scatter(cantilever_x_datas[k], cantilever_normalized_y_datas[k], s=100, marker='s', edgecolor='black', linewidth = 1, label = data_labels[k])
    ax.plot(cantilever_x_datas[k], cantilever_normalized_y_datas[k], linewidth = 1)
axes_font_size = 16
title_font_size = 16
legend_font_size = 14
font_name = 'Arial'
font_weight = 'normal'
ax.set_xlabel('aTC Concentration (\u03BCM)',fontsize = axes_font_size,fontweight = font_weight, fontname = font_name)
ax.set_ylabel('Normalized Response \n$\mathregular{[DNA]/[DNA]_{0}}$ or De-bending/De-$\mathregular{bending_{max}}$',fontsize = axes_font_size,fontweight = font_weight, fontname = font_name)
ax.set_xscale('log')
ax.set_ylim([0, 1.1])
ax.get_xaxis().set_major_formatter(matplotlib.ticker.FuncFormatter(lambda x, p: format(np.around(x,3), ',')))
legend_font = font_manager.FontProperties(family=font_name,
                       weight=font_weight,
                       style='normal', 
                        size=legend_font_size)
ticks_font = font_manager.FontProperties(family=font_name,
                   weight=font_weight,
                   style='normal', 
                    size=legend_font_size)
tkw = dict(size = 6, width = 1.5)
ax.tick_params(axis ='y',
         **tkw)
ax.tick_params(axis ='x',
         **tkw)
# Shrink current axis by 20%
box = ax.get_position()
ax.set_position([box.x0, box.y0, box.width * 0.8, box.height])
leg = ax.legend(frameon=False,prop=legend_font,loc='center left', bbox_to_anchor=(1, 0.5))
for label in ax.get_xticklabels():
    label.set_fontproperties(ticks_font)
for label in ax.get_yticklabels():
    label.set_fontproperties(ticks_font)
for line in leg.get_lines():
    line.set_linewidth(line_width_for_plot)
for axis in ['top', 'bottom', 'left', 'right']:
    ax.spines[axis].set_linewidth(1.5)  # change width
    ax.spines[axis].set_color('black')
ax.get_xaxis().set_major_formatter(matplotlib.ticker.FuncFormatter(lambda x, p: format(np.around(x,3), ',')))

plt.show()

    

# %% [code]


# %% [code]
```
